# KEK-6: A TRUNCATED-TRK-LIKE RECEPTOR FOR *DROSOPHILA* NEUROTROPHIN 2 REGULATES STRUCTURAL SYNAPTIC PLASTICITY

**DOI:** 10.1101/124644

**Authors:** Suzana Ulian-Benitez, Simon Bishop, Istvan Foldi, Jill Wentzell, Chinenye Okenwa, Manuel G. Forero, Bangfu Zhu, Marta Moreira, Mark Phizacklea, Graham McIlroy, Nicholas J Gay, Alicia Hidalgo

**Affiliations:** NeuroDevelopment Group, School of Biosciences, University of Birmingham, Edgbaston, Birmingham B15 2TT, United Kingdom.; Grupo D+Tec, Universidad de Ibagué, Colombia.; Department of Biochemistry, University of Cambridge, United Kingdom.

**Author notes:** These authors contributed equally (Suzana Ulian-Benitez, Simon Bishop). Author for correspondence, www.biosciences-labs.bham.ac.uk/hidalgo, http://www.birmingham.ac.uk/staff/profiles/biosciences/hidalgo-alicia.aspx.

**Keywords:** CNS, neuron, neuromuscular junction, *Drosophila*, neurotrophin, structural plasticity, structural homeostasis, synapse, DNT1, DNT2, receptor, Kekkon, Kek, Trk, TrkB-T1, Toll-6, CaMKII, VAP33A

## Abstract

Neurotrophism, structural plasticity, learning and long-term memory in mammals critically depend on neurotrophins binding Trk receptors to activate tyrosine kinase (TyrK) signalling, but *Drosophila* lacks full-length Trks, raising the question of how these processes occur in the fly. Paradoxically, truncated Trk isoforms lacking the TyrK predominate in the adult human brain, but whether they have neuronal functions independently of full-length Trks is unknown. *Drosophila* has TyrK-less Trk-family receptors, encoded by the *kekkon (kek)* genes, suggesting that evolutionarily conserved functions for this receptor class may exist. Here, we asked whether Keks function together with Drosophila neurotrophins (DNTs) at the larval glutamatergic neuromuscular junction (NMJ). Starting with an unbiased approach, we tested the evelen LRR and Ig-containing (LIG) proteins encoded in the *Drosophila* genome for expression in the central nervous system (CNS) and potential interaction with DNTs. Kek-6 was expressed in the CNS, could interact genetically with DNTs and could bind DNT2 both in signaling essays and in co-immunoprecipitations. There is promiscuity in ligand binding, as Kek-6 could also bind DNT1, and Kek-5 could also bind DNT2. In vivo, Kek-6 is found presynaptically in motoneurons, and binds DNT2 produced by the muscle, which functions as a retrograde factor at the NMJ. Kek-6 and DNT2 regulate NMJ growth, bouton formation and active zone homeostasis. Kek-6 does not antagonise the alternative DNT2 receptor Toll-6, but rather the two receptors contribute in distinct manners to NMJ structural plasticity. Using pull-down assays, we identified and validated CaMKII and VAP33A as intracellular partners of Kek-6, and show that together they regulate NMJ growth and active zone formation. These functions of Kek-6 could be evolutionarily conserved, raising the intriguing possibility that a novel mechanism of structural synaptic plasticity involving truncated Trk-family receptors independently of TyrK signaling may also operate in the human brain.

**AUTHOR SUMMARY:** A long-standing paradox had been to explain how brain structural plasticity, learning and long-term memory might occur in Drosophila in the absence of canonical Trk receptors for neurotrophin (NT) ligands. NTs link structure and function in the brain enabling adjustments in cell number, dendritic, axonal and synaptic patterns, in response to neuronal activity. These events are essential for brain development, learning and long-term memory, and are thought to depend on the tyrosine-kinase function of the NT Trk receptors. However, paradoxically, the most abundant Trk isoforms in the adult human brain lack the tyrosine kinase, and their neuronal function is unknown. Remarkably, Drosophila has kinase-less receptors of the Trk family encoded by the *kekkon (kek)* genes, suggesting that deep evolutionary functional conservation for this receptor class could be unveiled. Here, we show that Kek-6 is a receptor for Drosophila neurotrophin 2 (DNT2) that regulates structural synaptic plasticity via CaMKII and VAP33A, well-known factors regulating synaptic structure and plasticity, and vesicle release. Our findings suggest that in mammals truncated Trk-family receptors could also have synaptic functions in neurons independently of Tyrosine kinase signalling. This might reveal a novel mechanism of brain plasticity, with important implications for understanding also the human brain, in health and disease.

## INTRODUCTION

Brain plasticity, neurotrophism in development, structural and synaptic plasticity during learning and long-term memory in humans critically depend on the receptor TrkB binding its neurotrophin (NT) ligand BDNF [1,2]. During development, NTs and Trks regulate neuronal number and connectivity; subsequently BDNF and TrkB establish a reinforcing positive feedback loop promoting synaptic potentiation, and regulate the dynamic generation and elimination of synaptic boutons and dendritic spines in response to activity [1,3-5]. Thus, NTs and Trks are fundamental to linking structure and function in the brain, and this is thought to depend mostly on the tyrosine kinase (TyrK) function of Trks. Through its intracellular TyrK domain, TrkB activates the Ras/MAPKinase, PI3Kinase/ AKT and PLCγ signalling pathways downstream [1,2]. Pre-synaptic targets include Synapsin, to trigger vesicle release [5]. Post-synaptic targets include NMDAR and CREB, essential for long-term potentiation, learning and long-term memory [1,5]. Paradoxically, full-length TrkB decays postnatally and TrkB homodimers are not formed in the adult mammalian brain [6-10]. Instead, the most abundant adult isoform is the truncated TrkB-T1 isoform lacking the TyrK [8-10]. Mutant mice lacking TrkB-T1 have anxiety, and in humans alterations in TrkB-T1 are linked to severe mental health disorders [11-13]. However, the neuronal functions of the truncated Trk isoforms are poorly understood.

No canonical, bona fide, full-length, TyrK Trk receptors have been found in *Drosophila*. However, neurotrophism, structural and synaptic plasticity, learning and long-term memory all occur in fruit-flies, implying that either in the course of evolution insects and humans found different molecular solutions to elicit equivalent functions, or that undiscovered mechanisms contribute to brain plasticity in both humans and fruit-flies. Finding out what happened to the Trks in *Drosophila* is important, as it could uncover novel fundamental mechanisms of structure-function relationships in any brain.

Trk receptors have long been searched for in *Drosophila*. Trks (TrkA,B,C) bear the unique combination of Cysteine Rich Repeats (CRR), Leucine Rich Repeats (LRR) and Immunoglobulin (Ig) domains extracellularly, and an intracellular TyrK domain (Fig. 1A). Original searches focused on the TyrK domain, and identified DTrk and Dror as candidate *Drosophila* Trk homologues, but these are unlikely to bind neurotrophins. DTrk, also known as Off-Track (Otk)[14,15] lacks the LRR and CRR modules, it is kinase-dead and binds Semaphorins (Fig. 1A). Dror/Dnrk, like all Ror-family receptors, has an extracellular Frizzled/Kringle module instead [16-18](Fig. 1A). Subsequent proteomic analyses found no full-length, canonical Trk orthologues with the combination of LRR, CRR and Ig modules extracellularly and a Trk-family TyrK intracellularly in *Drosophila* [19-21]. However, phylogenetic analysis of the Trk-receptor superfamily identified the *Drosophila* Kekkons (Keks), lacking an intracellular TyrK domain, as closely related to the Trks [22](Fig. 1A,B). Trks and Keks both belong to the LIG family of proteins that contain extracellular ligand-binding LRR and Ig motifs [22,23]. There are 38 LIGs in humans, and amongst these are transmembrane proteins with a divergent intracellular domain lacking a TyrK or any conserved motifs [22]. There are 9 LIGs in *Drosophila* and phylogenetic analysis clusters mammalian AMIGO, LINGO, LRIG and LRRC4 in one clade with *Drosophila* Lambik, and mammalian Trks in a separate clade together with *Drosophila* Keks (Kek1-6) [22](Fig. 1B). Keks are more similar to the Trks than all other vertebrate LIGs are to each other [22]. Keks have only been found in insects, and thus are remnant, conserved Trk-like receptors in fruit-flies.

**Fig.1.**
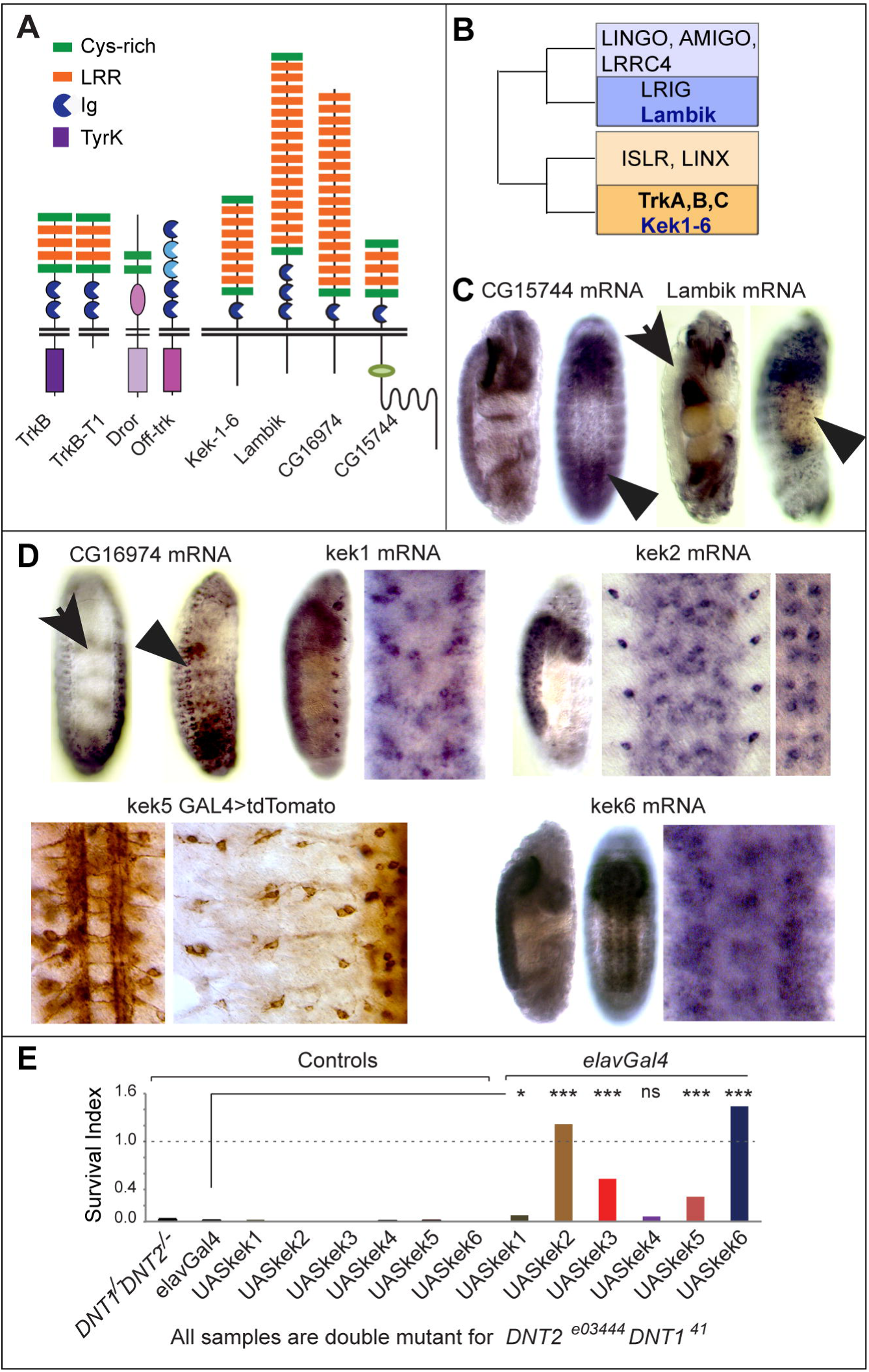
Keks are Trk-like receptors expressed in the CNS. (A) Modular composition of TrkB, TrkB-T1, Dror, Otk and *Drosophila* LIGs. (B) Amongst the LIGs, Keks are closer to the Trks than any other mammalian or Drosophila LIGs, adapted from the phylogeny of Mandai et al.[22]. (C-D) mRNA distribution in embryos: (C) *CG15744, lambik* and *CG16974* are not expressed in the VNC (arrows) above background, but *lambik* is in PNS and *CG16974* in muscle precursors (arrowheads); (D) *kek-1, kek-2* and *kek-6* transcripts are found in the VNC, and *kek5GAL4*>*tdTomato* drives expression in VNC and PNS (right) neurons. (E) Over-expression of *keks* – most prominently *kek2* and *6 -* in all neurons with *elavGAL4* rescued the cold semi-lethality of *DNT1^41^ DNT2^e03444^* double mutants. Chi-square and Bonferroni multiple comparisons correction. *p<0.05, ***p<0.001. For statistical details see Table S1.

There are 6 Keks in *Drosophila*, with a cytoplasmic tail lacking any remarkable conservation, except that Kek1, 2, 5 and 6 have a PDZ domain that could bind cytoplasmic effectors [24]. Kek1, 2, 5 and 6 are highly conserved in insects [24]. At least *kek1, kek2* and *kek5* are expressed in the CNS [25-27]. Only Kek2 has been investigated in the CNS, where it functions as a neuronal activity dependent modulator of synaptic growth and activity [27]. The ligands for the Keks have not been identified.

The prime candidates for Kek ligands are the *Drosophila* neurotrophins (DNTs). DNT1 (*Drosophila* neurotrophin 1 also known as Spz2), DNT2 (also known as Spz5) and Spz bear the distinctive and evolutionarily conserved neurotrophin cystine-knot domain of the mammalian neurotrophins [28-36], and they have conserved CNS functions regulating neuronal survival, connectivity and synaptogenesis [35-39]. Spz, DNT2 and DNT1 are known ligands for Toll-1, Toll-6 and Toll-7 receptors of the Toll and Toll-Like Receptor (TLR) superfamily [38,40]. In *Drosophila* Toll-1, Toll-6 and Toll-7 are required for targeting at the embryonic neuromuscular junction (NMJ), Toll-6 and Toll-8 for larval NMJ growth, and Toll-1, Toll-6 and Toll-7 function as neurotrophin receptors regulating neuronal survival and death, connectivity and behaviour [35-38,41-44]. In *Drosophila*, the NMJ is glutamatergic, and undergoes plasticity and potentiation, thus resembling mammalian central synapses, and is the standard context in which to investigate synaptic structural and functional plasticity [45]. Given that NT family ligands can bind multiple receptor types, and that receptors and ligands tend to co-evolve [17], conservation of the extracellular ligand-binding domain of Trks and Keks suggested Keks could potentially function as DNT receptors in flies.

Here, we investigated whether Kek-6 might function as a DNT receptor in *Drosophila*, at the glutamatergic NMJ synapse.

## RESULTS

### Kek6 is a truncated-Trk-like receptor that binds DNTs

To investigate if *Drosophila* LIGs might function as DNT receptors in the CNS, we first looked at their expression in embryos. *CG15744* did not reveal expression above background in the ventral nerve cord (VNC) (Fig. 1C); *lambik* and *CG16974* mRNAs were absent from the VNC, but might be expressed in the Peripheral Nervous System (PNS) or muscle, respectively (Fig. 1C,D). *kek1* and *kek2* were expressed in VNC cells, and possibly in PNS cells too (Fig. 1D), as previously shown [25,27]. *kek3* and *kek4* transcripts were not detected in embryos. *kek5* is expressed in the VNC [26], and we confirmed this with a *kek5-GAL4* reporter, which also revealed PNS expression (Fig. 1D). *kek6* transcripts were abundant in the VNC (Fig. 1D). Thus, amongst the 9 LIGs, Kek1, 2, 5, and 6 could function in the CNS.

To test whether Keks could function downstream of DNTs in vivo, we took advantage of the cold semi-lethality of *DNT1^41^ DNT2^e03444^* double mutants [38], and asked whether it could be rescued with the over-expression of *keks* in neurons. Over-expression of *kek1* and *kek4* in neurons (with *elavGAL4*) did not rescue, and *kek2* and *kek6* did most prominently (Fig. 1E, Table S1). Thus, Kek2 and Kek6 could function downstream of DNTs.

To ask whether DNT ligands could bind Keks and induce signaling, as Keks lack the TyrK, to enable a signaling readout, we generated chimaeric receptors formed of the extracellular and transmembrane domains of the Keks fused to the intracellular domain of Toll-6 (Fig. 2A). Toll-6 uses a conserved TIR domain to activate Dif/NFκB signaling downstream, which can be measured through the activation of *drosomicin-luciferase (dros-luc)* [38]. We used S2 cells stably transfected with *dros-luc*, transfected them with *kek-Toll-6* chimaeric receptors, and tested whether stimulation with purified cleaved DNT2 (DNT2-CK) could induce Dif/NFκB signaling. The chimaeric receptors targeted correctly to the S2 cell membrane (Fig. 2A). Stimulation with DNT2-CK in pDONR transfected controls induced *dros-luc*, as S2 cells express multiple Toll receptors [36,38] (Fig. 2A). Stimulation with DNT2-CK of cells transfected with *kek3*,*4*,*5*,*6-Toll-6* chimaeras had an effect comparable to the induction by stimulated full-length *Toll-6*, and Kek3-Toll-6 and Kek6-Toll-6 chimaeras responded more robustly (Fig. 2A, Table S1). We could not generate Kek1 and Kek2 chimeric receptors, thus a potential interaction with them cannot be ruled out. Thus, DNT2 can interact physically with at least Kek-3 and Kek-6. Together with the above data, this indicated Kek-6 was a strong candidate to interact with DNTs in the CNS.

**Fig.2.**
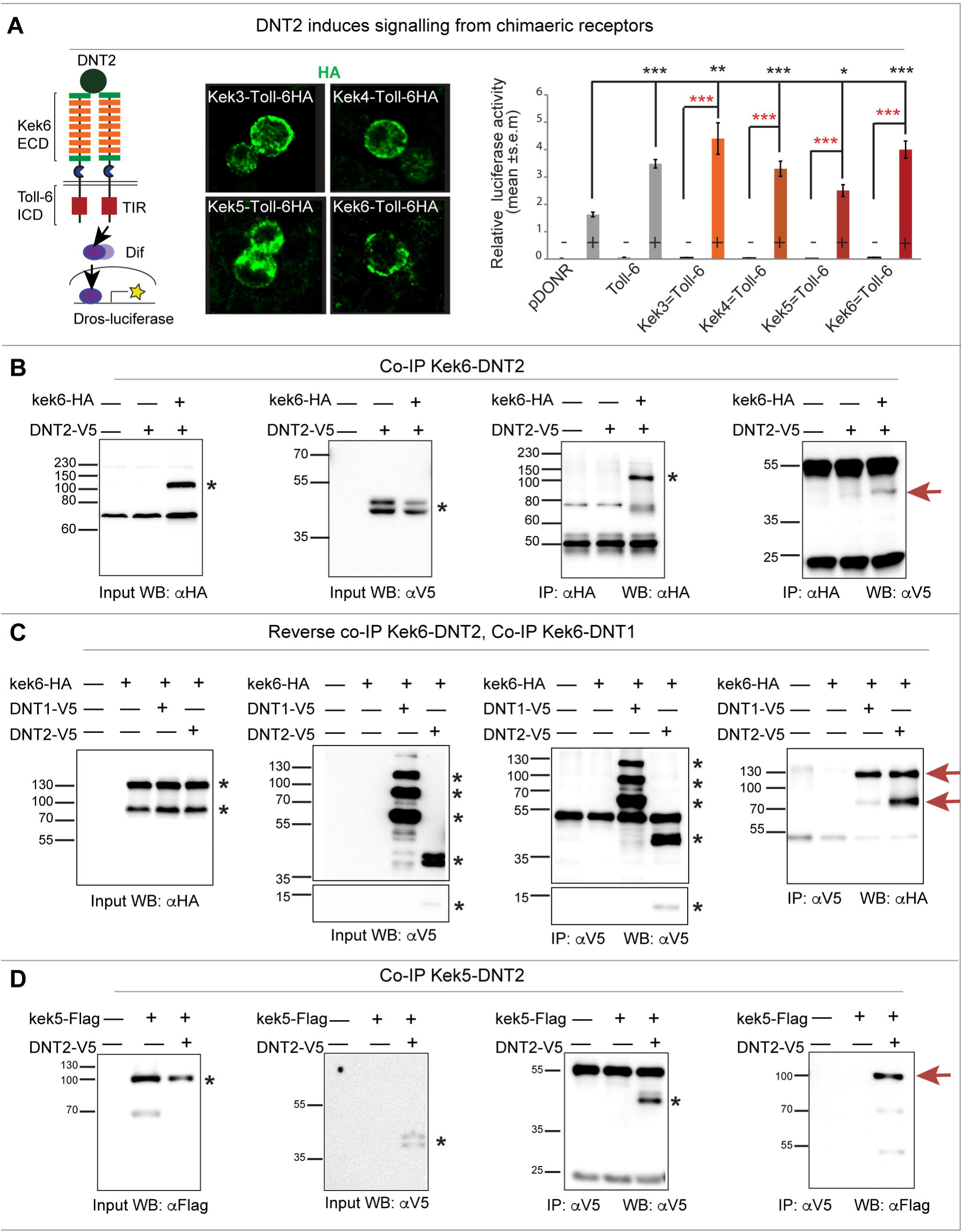
Keks bind DNTs. (A) Diagram of Chimaeric Kek-Toll-6-HA receptors, bearing the extracellular and transmemebrante domains of Keks and the intracellular signaling domain of Toll-6 (left). Chimaeric receptors visualized with anti-HA antibodies are distributed to the cell membrane in S2 cells (middle). Binding of purified mature DNT2-CK to the extracellular domain of Kek-Toll-6 receptors activates the *drosomycin-luciferase* reporter (right). Black asterisks denote comparisons to empty vector (pDONR) control, and red asterisks refer to no-stimulation controls. Welch ANOVA p<0.001; asterisks refer to post-hoc Games-Howell *p<0.05, **p<0.01, ***p<0.001. (B-D) Western blots showing co-immunoprecipitations from co-transfected S2 cells: (B) precipitation of Kek6-HA with anti-HA brings down DNT2-V5 detected with anti-V5. (C) Reverse co-IP for Kek6 and DNT2. Precipitation of DNT1-V5 also brings down Kek6-HA. (D) Precipitation of DNT2-V5 brings down Kek5-Flag. Black asterisks indicate the relevant control bands, red arrows indicate co-IPs. Input controls: cell lysate from co-transfected cells express both proteins. IP: immunoprecipitation; WB: western blot; FL: full length; CK: cystine-knot cleaved form. DNTs can be spontaneously cleaved following expression from these constructs in S2 cells.

To verify whether Kek-6 could bind DNT2, we carried out co-immunoprecipitations. S2 cells were co-transfected with HA-tagged *kek-6* and V5-tagged full-length *DNT2*. Precipitating Kek6-HA with anti-HA, brought down DNT2-V5 detected with anti-V5 (Fig. 2B). Conversely, precipitating DNT2 with anti-V5, also brought down Kek6 detected with anti-HA (Fig. 2C). Thus, Kek6 can bind DNT2. To test whether Kek6 might also bind DNT1, we co-transfected S2 cells with *kek6-HA* and full-length *DNT1-V5*. Precipitating DNT1 brought down Kek6 (Fig. 2C). Thus, Kek6 can bind both DNT2 and DNT1. To test whether another Kek might also bind the DNTs, S2 cells were co-transfected with *kek5-Flag* and *DNT2-V5*, and we found that precipitating DNT2 also brought down Kek-5 (Fig. 2D). Therefore, Kek-5 can also bind DNT2. These data demonstrate that Keks bind DNT ligands, and that binding is promiscuous.

To conclude, Kek-6 was widely expressed in the CNS, rescued the semi-lethality of *DNT1 DNT2* double mutants, and bound DNT ligands in a signaling assay and in co-immunoprecipitations. Thus, Kek-6 could function as a DNT receptor.

### Kek-6 in motoneurons and DNT2 from the muscle function together at the NMJ

For a functional *in vivo* analysis we focused on Kek-6. *kek-6* CNS expression was examined in larvae using Kek6^MIMIC13953^ flies bearing a GFP insertion into the *kek-6* coding region (hereafter named Kek6^GFP^). Kek6^GFP^ was found in Eve+(Fig. 3A) and HB9+(Fig. 3B) interneurons and motoneurons, and excluded from Repo+ glia (Supplemental Fig.S1) in third instar larval VNCs. Kek6^GFP^ was present in motoneuron terminals at the neuro muscular junction (NMJ) of third instar larvae and not in the muscle (Fig. 3C,D), revealing NMJ6/7 synaptic boutons, surrounded by post-synaptic Dlg (i.e. *Drosophila* PSD95) (Fig. 3D,E). Thus, *kek-6* is expressed pre-synaptically in motoneurons.

**Fig.3.**
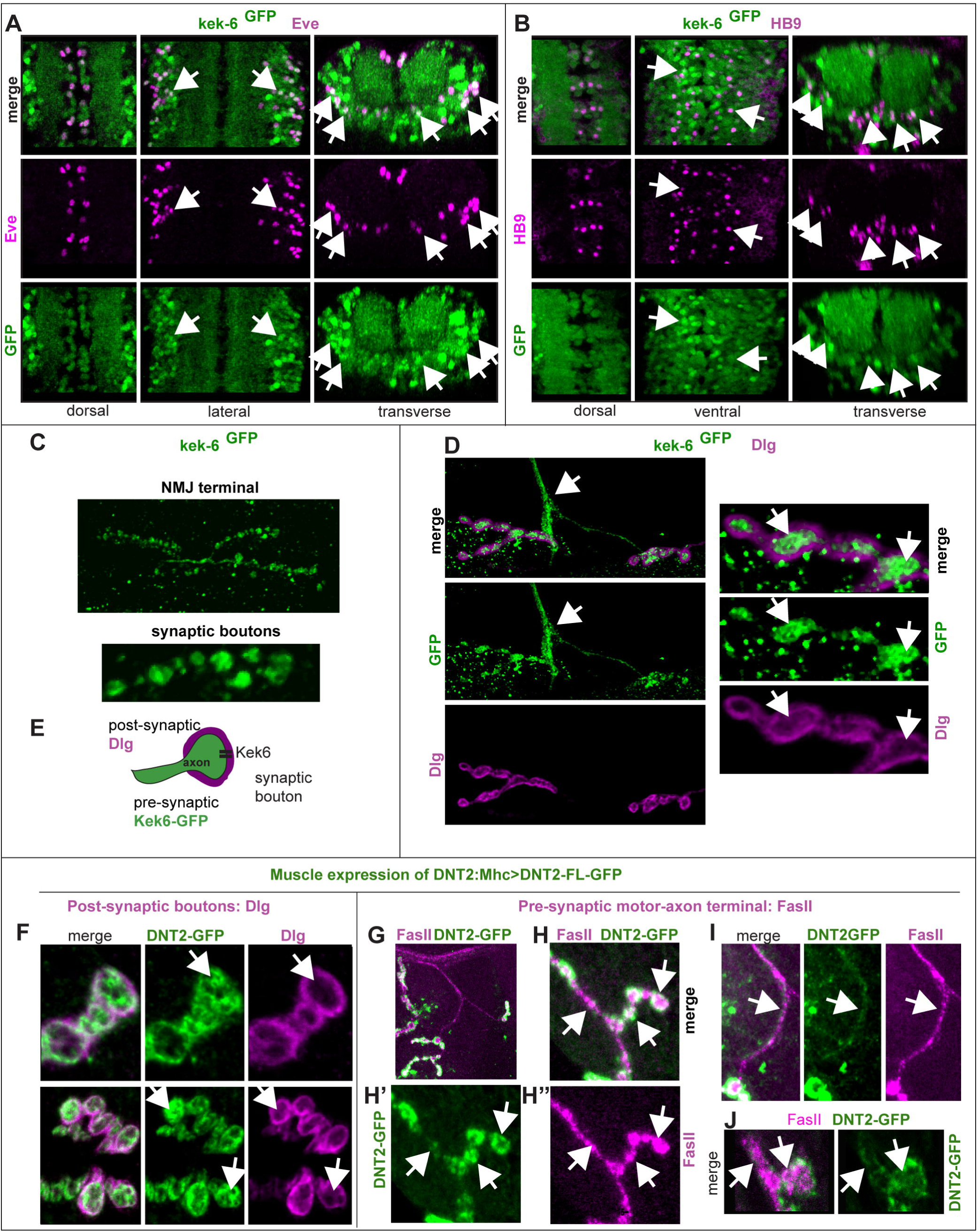
Kek-6 is expressed pre-synaptically in motoneurons and binds post-synaptic DNT2. (A-B) Kek-6^GFP^ GFP colocalises with the neuronal markers Eve (A) and HB9 (B) in the larval VNC (arrows show examples). (C) Kek-6^GFP^ was found in third instar larval muscle 6/7 NMJ and synaptic boutons (higher magnification). (D) Kek-6^GFP^ was found in the motorneuron axonal terminal (arrows), and in pre-synaptic bouton lumen (higher magnification on the right), not colocalising with the post-synaptic marker anti-Dlg (arrows). (E) Illustration. (F-J) Over-expression of GFP tagged full-length DNT2 in muscle (*MhcGAL4*>*UAS-DNT2-FL-GFP*) revealed: (F) DNT2-GFP distribution within the pre-synaptic bouton lumen (arrows), boutons labeled post-synaptically with anti-Dlg; (G-J) DNT2-GFP along the motoraxon and within the pre-synaptic bouton lumen, both labeled with FasII (arrows).

*DNT2* transcripts are expressed in the larval body wall muscles, and localised to post-synaptic boutons [39]. To test if DNT2 could function retrogradely, we generated a tagged form of full-length DNT2 with GFP at the C-terminus, and over-expressed *DNT2-FL-GFP* in the muscle with *MhcGAL4*. DNT2-FL-GFP would result in secretion of mature DNT2-CK-GFP [36], although not all protein might get cleaved and secreted. Over-expression of *DNT2-FL-GFP* in muscle resulted in the localization of GFP pre-synaptically in boutons, surrounded by the post-synaptic marker Dlg (Fig. 3F). DNT2-FL-GFP also colocalised with the motoneuron marker FasII in boutons (Fig. 3G,H,J) and in motoraxons (Fig. 3H,I,J). Thus, DNT2 produced in muscle can interact with the receptor Kek-6 in motoneurons, consistent with a retrograde function.

To investigate the *in vivo* functions of Kek6 and DNT2, we generated *kek6* and *DNT2* null mutant alleles by FRT-mediated recombination between PiggyBac insertions [46]. Neither *kek6^34^/Df(3R)ED6361* or *DNT2^37^/Df(3L)6092* loss of function mutants, nor *kek6^−/−^ DNT2^−/−^* double mutants, affected viability. We analysed larval locomotion using FlyTracker software to trace trajectories and measure crawling speed [38], and found that *kek6^34^/Df(3R)ED6361* mutant larvae crawled more slowly than controls (Fig. 4A, Table S1), and *DNT2^37^/Df(3L)6092* larvae crawled even slower (Fig. 4A, Table S1). *kek6^−/−^ DNT2^−/−^*double null larvae crawled at similar speeds as *DNT2^−/−^* mutants, but travelled shorter distances, moving around the starting spot (Fig. 4A, Table S1). The synergistic effect in the double mutants suggested that Kek-6 and DNT2 are functionally linked.

**Fig. 4.**
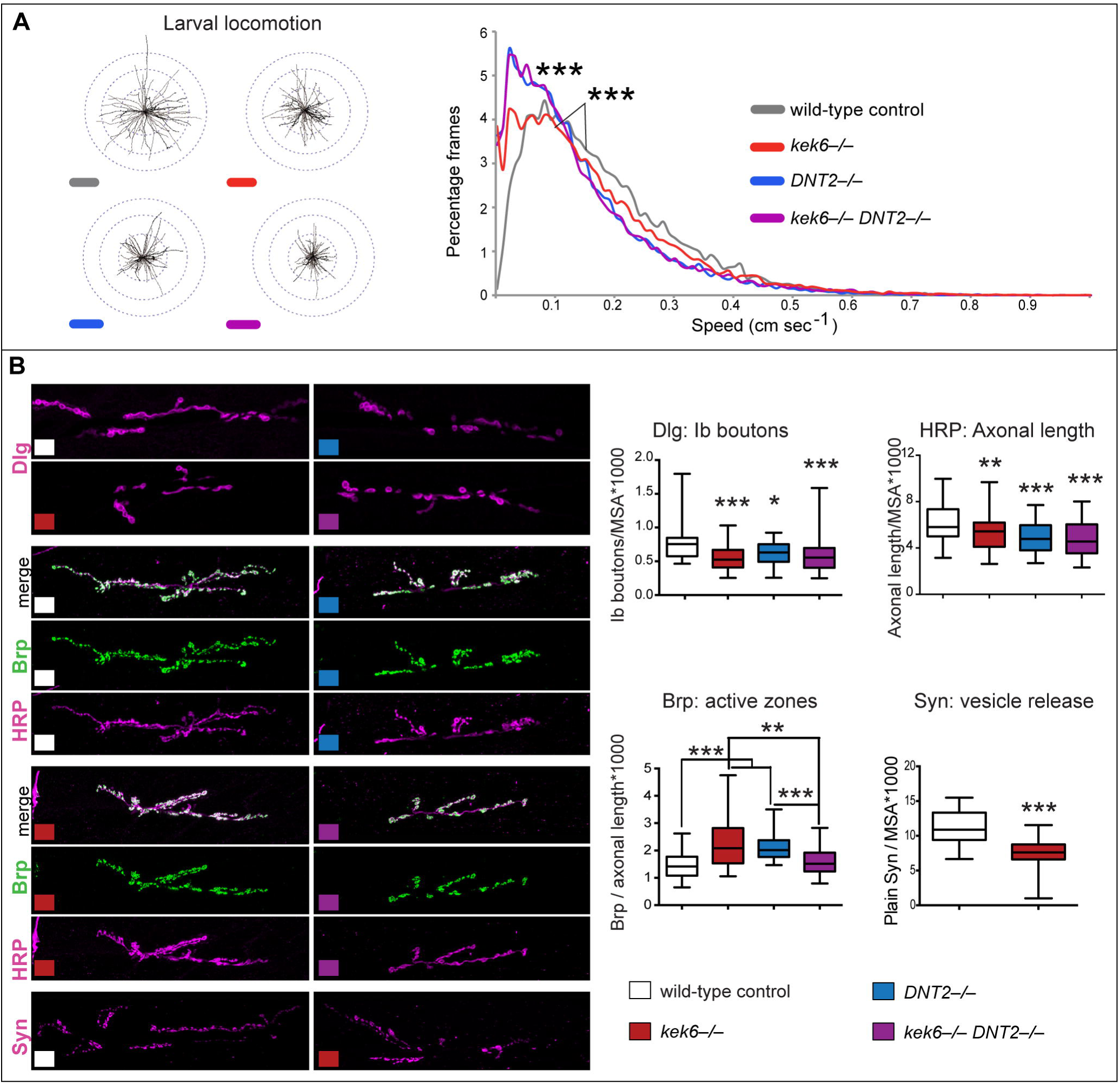
*kek6* and *DNT2* mutants have smaller NMJs. (A) Plotted trajectories of filmed larvae (left), and histograms (right) of frames vs. speed analysed with FlyTracker. Kruskal-Wallis p<0.0001 and ***p<0.001 post-hoc Dunn test. (B) NMJs (left) and box-plot graphs (right) showing: *kek-6^−/−^* and *DNT2^−/−^* single mutants and *kek-6^−/−^ DNT2^−/−^* double mutants have fewer Dlg+ boutons and smaller HRP+ axonal terminals (normalized to muscle area, MSA). Dlg: Kruskal-Wallis p<0.0001, and *p<0.05, **p<0.01, ***p<0.001 post-hoc Dunn; HRP: One Way ANOVA p<0.0001, and **p<0.01, ***p<0.001 post-hoc Dunnett. *kek-6* and *DNT2* single mutants, but not the double mutants, have increased active zone density (Brp+/HRP+axonal length). Brp: Kruskal-Wallis p=0.0012, and **p<0.01, ***p<0.001 Dunn’s post-hoc. *Kek-6* mutants have reduced Synapsin, Mann-Whitney U test ***p<0.001. For statistical details, see Table S1. Mutant genotypes throughout figures: Control: *yw*/+; Mutants: *kek-6^−/−^ kek6^34^/Df(3R)6361; DNT2^−/−^ : DNT2^37^/Df(3L)6092; kek-6^−/−^ DNT2^−/−^ : kek6^34^Df(3L)6092/ Df(3R)6361 DNT2^37^*.

Locomotion phenotypes suggested the NMJ might be affected, so we looked at the muscle 6/7 NMJs, which require DNT1 and 2 [39]. Targeting at muscle 6/7 NMJ was affected in *kek6^−/−^* mutant embryos and upon *kek-6* over-expression in neurons (Supplemental Fig.S2). In wandering third instar larvae, *kek6^−/−^* and *DNT2^−/−^* single mutant larvae, and *kek6^−/−^ DNT2^−/−^* double mutants, had smaller NMJs than controls, with fewer Ib boutons and shorter axonal terminal length (Fig. 4B, Table S1). Thus, Kek-6 and DNT2 are required for normal NMJ growth. *kek6^−/−^* and *DNT2^−/−^* single mutant NMJs had higher active zone density, visualized with anti-Brp (*Drosophila* ELKs) and quantified automatically throughout the NMJ stack of images using DeadEasy Synapse software [39](Fig. 4B, Table S1). Since the NMJs were smaller, this suggested that the increase in active zones was a homeostatic compensation of defective synaptic function, to enable adequate behaviour. Homeostatic adjustments in active zones are a common manifestation of structural plasticity at the NMJ [47]. Remarkably, the increase in active zone density did not occur in *kek-6^−/−^ DNT2^−/−^* double mutants (Fig. 4B, Table S1), meaning that compensation fails in the double mutants. To further test how Kek-6 affects the synapse, we also visualised Synapsin, which phosphorylates components of the SNARE complex to promote neurotransmitter vesicle release [48], and quantified it automatically using DeadEasy Synapse software [39]. *kek6^−/−^* mutants had reduced Synapsin production (Fig. 4B, Table S1), revealing defective synapse composition. Conversely, over-expression of *kek-6* did not alter NMJ size (Fig. 5A and Supplemental Fig.S4A), but it increased active zone density (Fig. 5A, Table S1), and induced ghost boutons, albeit not significantly (Supplemental Fig.S3C-E, Table S1). Ghost boutons are presynaptic, HRP+ Dlg-negative, immature boutons that fail to get stabilized, and are correlates of increased neuronal activity, thus further indicating that Kek-6 influences structure and/or function. Like Kek-6 gain of function, over-expression of full-length *DNT2-FL* in muscle, and either full-length *DNT2-FL* or mature *DNT2-CK* in motorneurons (with *D42GAL4*), also increased active zones (Fig. 5B,C Table S1). Thus both DNT2 and Kek6 induce active zone formation. Altogether, these data show that both Kek-6 and DNT2 are required to regulate NMJ growth and for appropriate synaptic structure.

**Fig. 5.**
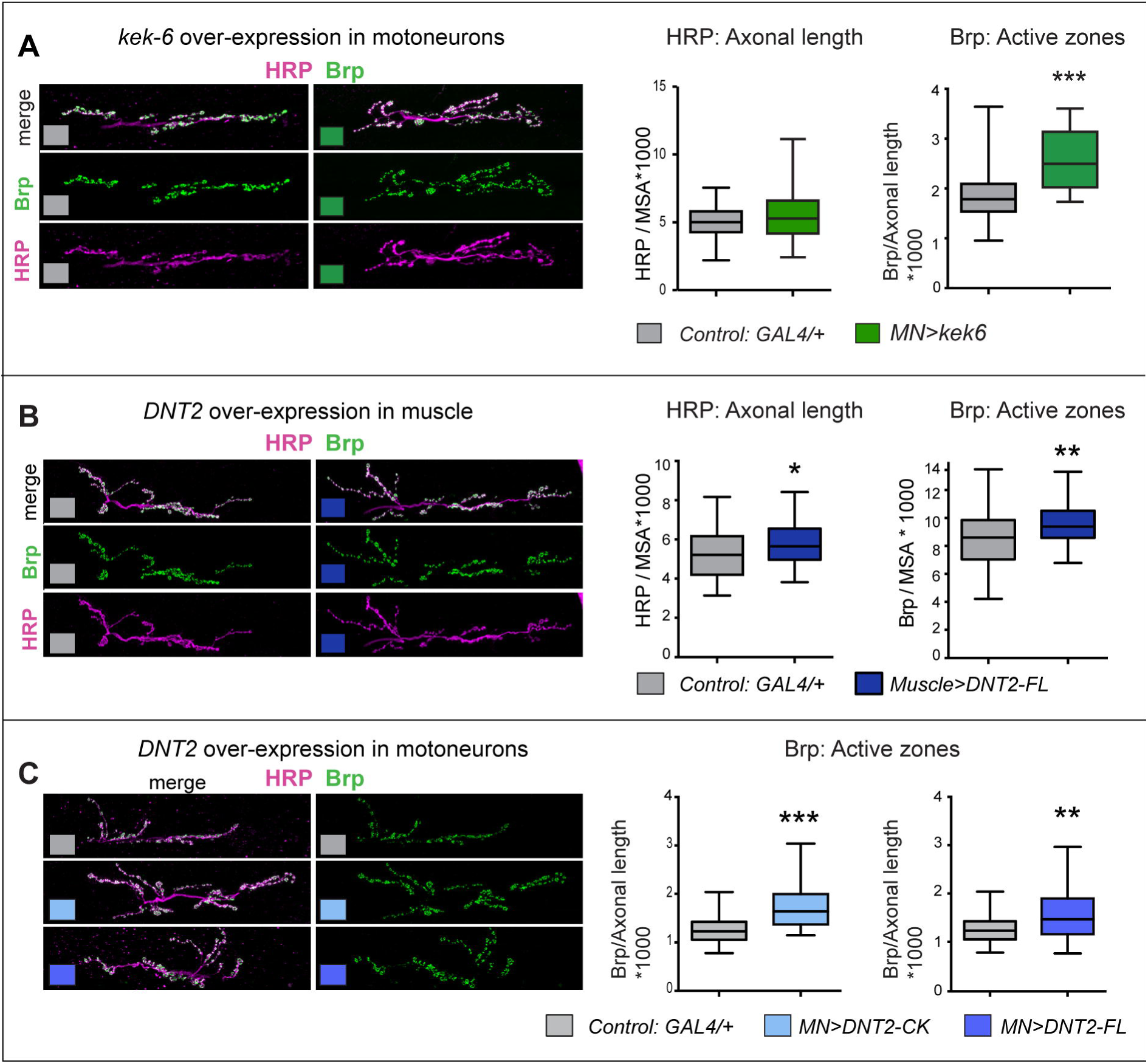
Kek-6 and DNT2 can induce active zones and NMJ growth. Confocal images of NMJs from A3-4 muscle 6/7 (left), and box-plot graphs (right), showing: (A) Over-expression of *kek6* in motoneurons had not effect on HRP+ NMJ size, but it increased Brp+ active zones. HRP: Student t test n.s. p=0.07; Brp: Mann-Whitney U test ***p<0.001.(B) Over-expression of full-length DNT2 in muscle increased NMJ size (HRP) and active zones (Brp), revealing a retrograde function. HRP: Mann-Whitney U test *p<0.05; Brp: Student t test **p<0.01. (C) Over-expression of both full-length DNT2 and mature DNT2-CK in motoneurons induced active zone formation. Brp DNT2-CK: Student t test **p<0.01, and Brp DNT2-FL: Mann-Whitney U test ***p<0.001. See Table S1. *MN*=*D42GAL4*.

Our data suggested that DNT2 functions as a retrograde ligand for Kek-6. Over-expression of *DNT2-FL* in muscle increased axonal terminal size and pre-synaptic active zones (Fig. 5B). To verify this, we used epistasis analysis. Over-expression of *kek6* in motorneurons rescued the NMJ mutant phenotypes of bouton number and axonal length of *kek6^34^/Df(3R)6361* mutants (Fig. 6A, Table S1), demonstrating that the *kek6-/-* mutant phenotypes were specific. Over-expression of *kek6* in motorneurons rescued the NMJ phenotypes of *DNT2^37^/Df(3L)6092* mutants and *kek6^−/−^ DNT2^−/−^* double mutants (Fig. 6B,C, Table S1), demonstrating that Kek-6 functions downstream of DNT2. Over-expression of untagged *DNT2-FL* in muscle (with *MhcGAL4*) increased Dlg+ bouton number (Fig. 6B), and importantly, this was rescued by *kek-6* loss of function (Fig. 6B, Table S1). Since Kek6 functions pre-synaptically, and DNT2 was over-expressed in muscle, this demonstrates that DNT2 is a retrograde ligand for Kek-6. Altogether, our data demonstrate that DNT2 is a retrograde ligand for Kek-6 at the NMJ and that DNT2 and Kek6 are required together for synaptic structure and NMJ growth.

**Fig. 6.**
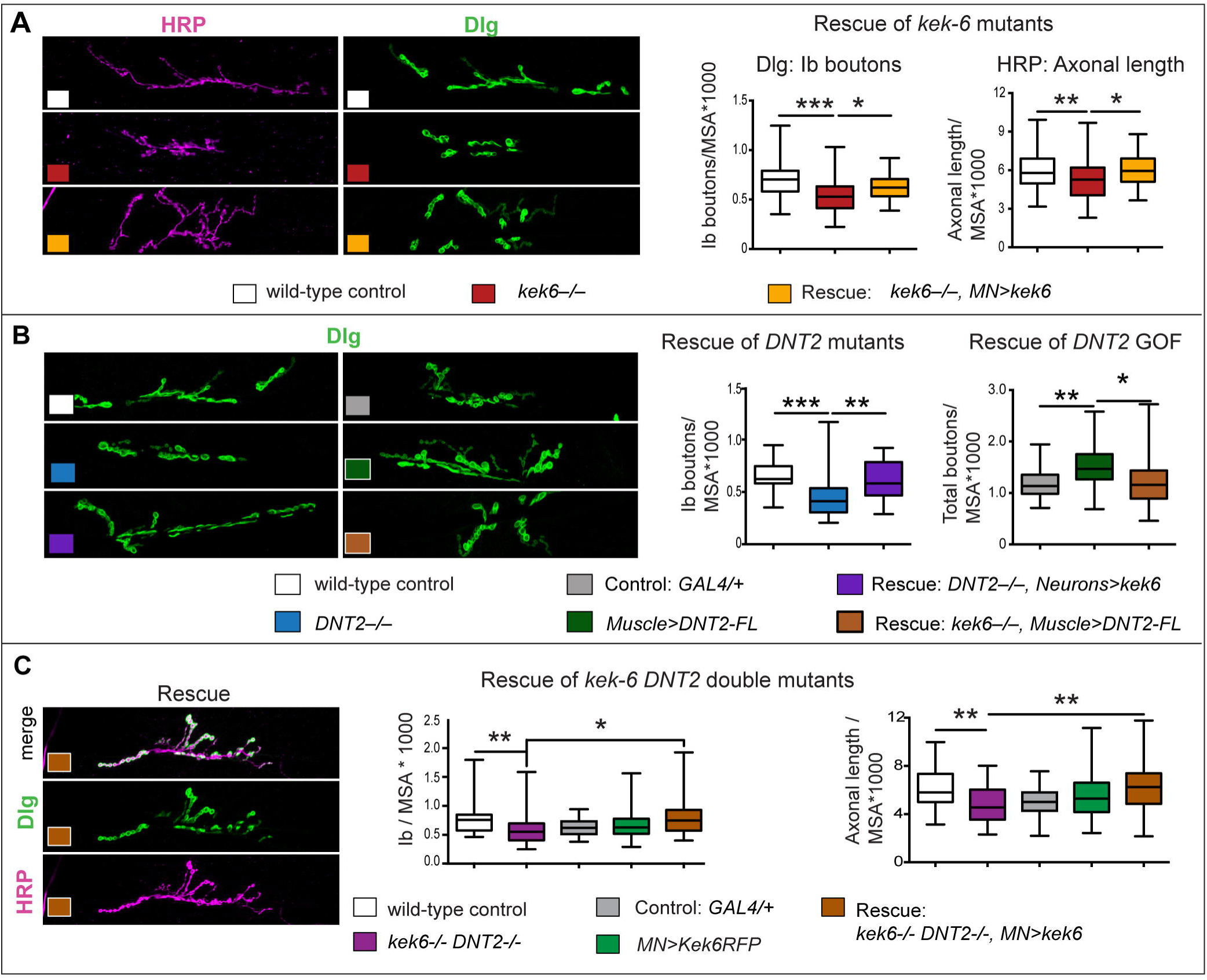
Kek-6 functions downstream of DNT2. Confocal images of NMJs from A3-4 muscle 6/7 (left), and box-plot graphs (right), showing: (A) Over-expression of *kek-6* in motoneurons rescued the phenotypes of *kek-6* mutants. Dlg: One Way ANOVA p<0.0001 and post-hoc Bonferroni *p<0.05, ***p<0.001. HRP: One Way ANOVA p<0.001 and post-hoc Bonferroni **p<0.01, **p<0.01. (B) Left: Over-expression of *kek-6* in neurons rescued the phenotype of *DNT2* mutants. Dlg: Kruskal-Wallis p=0.001, and post-hoc Dunn test **p<0.01, ***p<0.001. Right: *kek-6* loss of function rescued the increase in boutons caused by the muscle over-expression of *DNT2*. Dlg: Welch ANOVA p<0.01 and post-hoc Games-Howell *p<0.05, **p<0.01. (C) Over-expression of *kek-6* in motoneurons rescued the phenotypes of *kek-6 DNT-2* double mutants. Dlg: Kruskal-Wallis p=0.001 and post-hoc Dunn’s test *p<0.05, **p<0.01. HRP: Welch ANOVA p=0.000, post-hoc Games Howell **p<0.01. See Table S1. GAL4 drivers: Muscle: *MhcGAL4;* Neurons: *elavGAL4;* MN: *D42 or Toll-7GAL4*. Controls: white boxes: yw/+; grey boxes: GAL4/+; mutant genotypes as in Fig. 4. Rescue genotypes: (A) *w; UASkek6RFP/+; Df(3R)6361/kek6^34^D42GAL4*. (B) *w; UASkek6RFP/+; elavGAL4 Df(3L)6092/DNT2^37^*; and *w; UASDNT2-FL/+; Df(3R)6361/kek6^34^D42GAL4*. (C) *w; Toll-7GAL4/UASkek6RFP; kek6^34^Df(3L)6092/Df(3R)6361 DNT2^37^*.

Intriguingly, our data also suggested that DNT2 had additional functions compared to *kek-6*. Homeostatic compensation of active zones did not occur in *kek-6^−/−^ DNT2^−/−^* double mutants (Fig. 4B). Furthermore, over-expression of *DNT2* but not *kek-6* increased NMJ size, as both HRP+ axonal terminal length (Fig. 5B) and bouton number (Fig. 6B) increased when *DNT2-FL* was over-expressed from muscle. DNT2 is also a known ligand of Toll-6 [38], and Toll-6 and -8 are required in motorneurons for NMJ growth and active zone formation[37,43]. So this raised two questions: do Toll-6 and Kek-6 interact functionally as DNT2 receptors at the NMJ, and why?

### Kek-6 and Toll-6 cooperate to modulate NMJ structural homeostasis

LIG proteins and truncated Trk isoforms can function as ligand sinks or dominant negative co-receptors that abrogate signaling, e.g. by full-length Trks [6,7]. Thus, Kek-6 might inhibit Toll-6 function.

Although Toll-6 has NMJ functions, its expression here had not been reported [43]. Using a MIMIC insertion into the intronless coding region of Toll-6, we generated *Toll-6GAL4* flies by RMCE, and found Toll-6>mCD8-GFP to be distributed at the muscle 6/7 NMJ, colocalising with the motoraxon marker FasII (Fig. 7A). Like *kek-6* and *DNT2* mutants, *Toll-6^MIO2127^/Df(3L)BSC578* mutants had smaller NMJs, with shorter HRP+ axonal length (Fig. 6B, Table S1) as previously reported [43], confirming that Toll-6 is required for NMJ growth. Contrary to *kek-6^−/−^* mutants, *Toll-6^−/−^* mutants had reduced Brp+ active zones compared to controls (Fig. 7B), meaning that Toll-6 is required for active zone formation. *kek-6^−/−^ Toll-6^−/−^* double mutant larvae still had small NMJs, like each single mutant alone (Fig. 7B, Table S1), and Brp+ active zones remained lower than in controls and comparable to the levels of *Toll-6^−/−^* single mutants (Fig. 7B). These results reveal that: 1) Since Toll-6 is required for active zone formation, and over-expression of *kek-6* increased active zones, Kek-6 does not function as a DNT2 sink nor as an inhibitor or dominant negative co-receptor for Toll-6. 2) The compensatory increase in active zones observed in *kek-6^−/−^* and *DNT2^−/−^* single mutants did not occur in *kek-6^−/−^ DNT2^−/−^* or *kek-6^−/−^ Toll-6^−/−^* double mutants, meaning that the concerted functions of Toll-6 and Kek-6 as DNT2 receptors are required for NMJ structural homeostasis.

**Fig.7.**
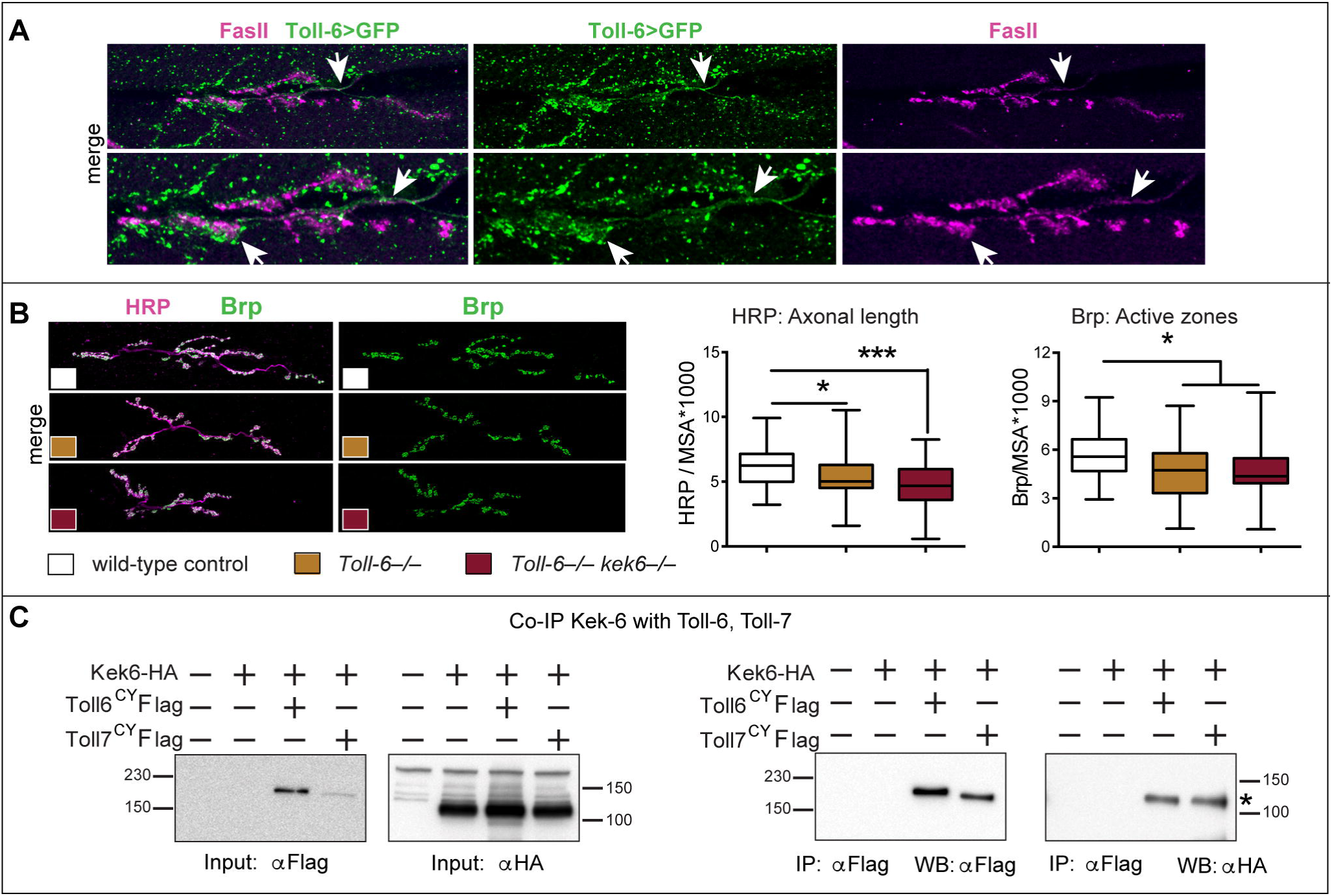
Kek-6 and Toll-6 interact for NMJ structural homeostasis. (A) *Toll-6GAL4*>*mCD8-GFP* revealed expression in motorneurons at the muscle 6/7 NMJ, colocalising with FasII presynaptically. (B) Muscle 6/7 NMJs (left) and box-plot graphs (right) showing: *Toll-6^MIO2127^/Df(3L)BSC578* mutants and *Toll-6^MIO2127^Df(3R)6361/kek6^35^ Df(3L)BSC578* double mutants had smaller NMJs (HRP, Kruskal-Wallis p=0.0001), and reduced active zones (Brp, Kruskal-Wallis p=0.0055), post-hoc Dunn for both *p<0.05, ***p<0.001. (C) Co-immunoprecitation from co-transfected S2 cells: Precipitating Toll-6 and Toll-7 with anti-Flag brought down Kek-6 detected with anti-HA. IP: immuno-precipitation; WB: western blot; asterisk: co-IP. See Table S1.

To test whether Kek-6 and Toll-6 might physically interact to form a receptor complex, we carried out co-immunoprecipitations, and to ask if potential interactions were promiscuous or specific, we also tested Toll-7. S2 cells were co-transfected with the active forms *Toll-6^CY^-Flag* or *Toll-7^CY^-Flag* and *Kek6-HA*, and we found that precipitating Toll-6^CY^ and Toll-7^CY^ with anti-Flag brought down Kek6 detected with anti-HA (Fig. 7C). Thus, Kek6 can bind Toll-6 and Toll-7.

Altogether, these data showed that retrograde DNT2 can bind two receptor types at the NMJ - Toll-6 and Kek-6 - which can interact pre-synaptically to cooperatively promote NMJ growth and regulate active zone homeostasis. This, however, raised a new question: if Kek-6 did not function simply by modulating Toll-6, and in the absence of a TyrK, how might it function?

### Kek-6 functions via CaMKII and VAP33A

To find out how Kek-6 might function, we carried out pull-down assays to isolate candidate factors binding its intracellular domain. S2 cells were transfected with *kek6-Flag*, and anti-Flag coated beads were used to expose Kek-6 to cell lysates from either S2 cells or wild-type adult fly heads, and bound proteins were isolated by SDS-PAGE followed by mass spectrometry (Fig. 8A,B). Candidates were identified as proteins present in Kek6-Flag samples and absent from non-transfected mock controls, and if identified from multiple peptides (Tables S2 and S3). Prevalent amongst these were proteins involved in vesicle trafficking, axonogenesis, dendrite morphogenesis and synaptic function (Fig. 8C). Amongst the top hits were CaMKinase II identified from fly heads, and VAP33A from both S2 cells and heads (Fig. 8C). CaMKII functions both as a kinase and a scaffolding protein, to promote structural synaptic plasticity. Post-synaptically, it phosphorylates and recruits AMPAR and NMDAR to the post-synaptic density, leading to post-synaptic potentiation, and pre-synaptically it localizes to active zones and phosphorylates Synapsin and other SNARE complex proteins, triggering neurotransmitter release [48-51]. VAP33A is a Vamp Associated Protein, of the SNARE complex, and also has universal functions in exocytosis and vesicle trafficking [52]. To validate these two candidates as downstream effectors of Kek-6, we carried out co-immunoprecipitations. We co-transfected S2 cells with Kek6-Flag and either CaMKII or VAP33A tagged with HA. Precipitating Kek-6-Flag, brought down CaMKII-HA (Fig. 8D). Similarly, precipitating Kek-6-Flag also brought down VAP33A-HA (Fig. 8D). Thus, Kek-6 can bind CaMKII and VAP33A.

**Fig.8.**
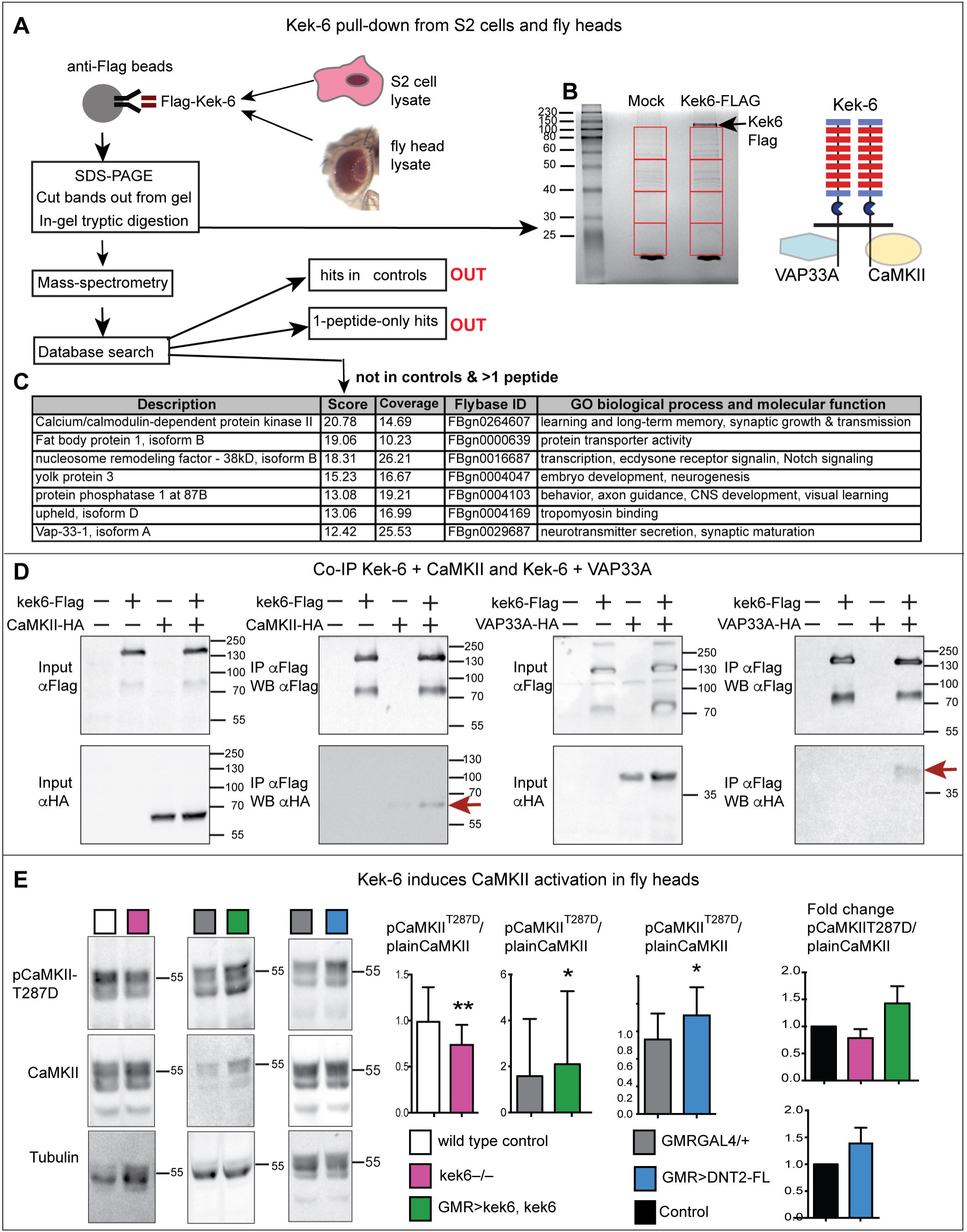
Kek-6 physically interacts with synaptic factors. (A) Diagram of pull-down workflow and (B) coomassie-stained gel showing proteins from S2 cell or adult fly head lysate bound to Kek-6-Flag or mock control. Red boxes indicate fractions used for mass spectrometry. Hits that appeared in mock controls, or with only one peptide, were disregarded. (C) Selection of candidate interacting proteins, see Tables S2 and S3. (D) Co-immunoprecipitations from co-transfected S2 cells: precipitating Kek-6-Flag brought down CaMKII-HA and VAP33A-HA. IP: immunoprecipitation, WB: western blot. (E) Western blots with anti-activated pCaMKII^T287^, non-phosphorylated plain CaMKII and Tubulin, from adult fly heads, GMRGAL4 drives expression in retina. Graphs show pCaMKII^T287^ levels normalized over non-phosphorylated CaMKII. Student paired t-tests. *p<0.05, **p<0.01, see Table S1.

To validate the functional relationship between Kek-6 and CaMKII *in vivo*, we asked whether altering Kek-6 function would affect CaMKII activation in brain. We tested for the constitutively active state of CaMKII, corresponding to phosphorylation at Thr286 (T287 in *Drosophila*), using anti-pCaMKII^T287^ antibodies. In the heads of *kek6^34^/Df(3R)ED6361* mutant adult flies the relative levels of pCaMKII^T287^, normalised over non-phosphorylated CaMKII, decreased (Fig. 8E Table S1). Conversely, over-expression of *kek-6* in retina (with *GMRGAL4*) increased CaMKII phosphorylation (Fig. 8E, Table S1). Over-expressing *DNT2-FL* had the same effect (Fig. 8E). Thus, Kek-6 is required for CaMKII activation, and both Kek-6 and DNT2 can activate CaMKII downstream.

Next we asked whether CaMKII and VAP33A might function downstream of Kek6 at the NMJ. Unfortunately, we were not able to get antibodies to inactive CaMKII to work reliably at the NMJ for normalisation, so we visualised constitutively activated CaMKII with pCaMKII^T287^ and quantified it automatically, using DeadEasy Synapse, which detected the increase in pCaMKII caused by the over-expression of activated CaMKII^T287^ in motoneurons (*D42*>*CaMKII^T287D^*, Fig. S4A,C). Pre-synaptic over-expression of *CaMKII^T287^* increased axonal length, Ib bouton number and active zone density, and inhibiting CaMKII activation pre-synaptically by over-expressing the CaMKII phosphorylation inhibitor *Ala* [53] resulted in smaller NMJs and reduced active zones (Fig.S4A-C, Table S1), as previously reported [54]. Thus we asked whether Kek-6 influenced CaMKII at the NMJ. Pre-synaptic over-expression of *kek6* increased pCaMKII^T287^ levels (Fig. 9A, Table S1). This increase was rescued by the over-expression of *Ala* together with *kek6*, showing that this phenotype was specific (Fig. 9A, Table S1). However, over-expression of *Ala* alone did not result in a detectable reduction in pCaMKII in this case (Fig. 9A, Table S1). This could be due either to the fact that CaMKII is most abundant post-synaptically, and we were only inhibiting it pre-synaptically, or to limited efficacy of the inhibitor. To try an alternative approach, we knocked-down pre-synaptic *CaMKII* expression with RNAi, and this decreased pCaMKII^T287^ levels, causing a stronger effect than Ala (Fig. 9A, Table S1). Like Ala, pre-synaptic *CaMKII-RNAi* knock-down together with *kek-6* over-expression also rescued pCaMKII^T287^ levels, and reduced them further than controls (Fig. 9A, Table S1), meaning that CaMKII functions downstream of Kek-6. Together, these data show that Kek-6 activates CaMKII.

**Fig. 9.**
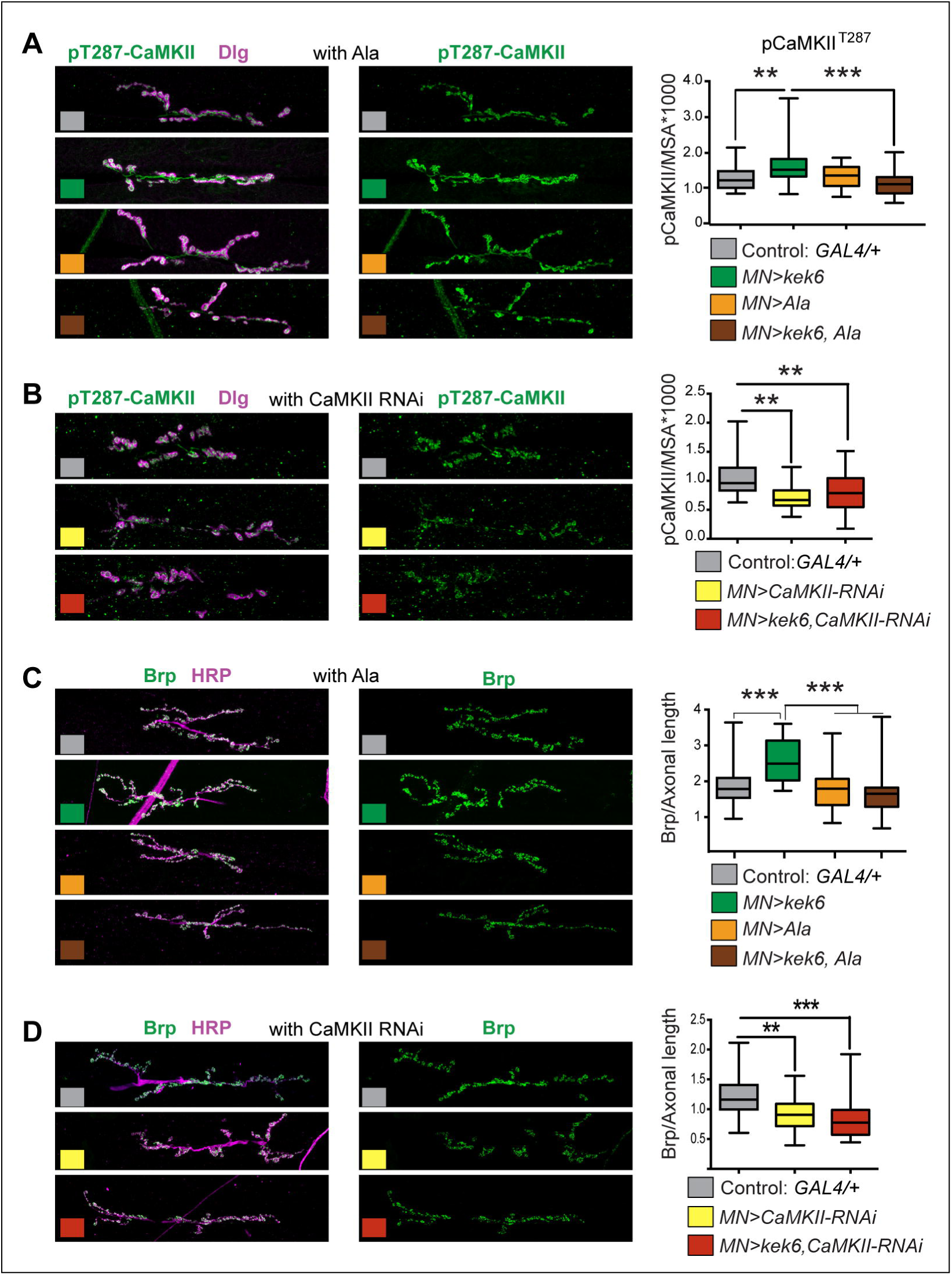
Kek6 activates CaMKII at the NMJ. NMJs from A3-4 muscle 6/7 (left), and box-plot graphs (right), NMJs, labeled with anti-pCaMKII^T287^ for the constitutively active form; anti-Dlg for post-synaptic boutons; anti-HRP for pre-synaptic axonal terminal length; anti-Brp for active zones. Brp and pCaMKII^T287^ were quantified automatically with DeadEasy Synapse. (A, B) Over-expression of *kek-6* in motoneurons increased pCaMKII^T287^ levels, which was rescued with the over-expression of the CaMKII inhibitor Ala (A) *or* CaMKII RNAi knock-down (B) pre-synaptically. Kruskal-Wallis p<0.0001 and **p<0.01, ***p<0.001post-hoc Dunn for both graphs. (C,D) CaMKII inhibition with Ala (C) or knock-down with CaMKII-RNAi (D) rescued the increase in active zones caused by *kek-6* over-expression. (C) Welch ANOVA p<0.000 and ***p<0.001 post-hoc Games-Howell, (D) Kurskal-Wallis p<0.0001 and **p<0.01, ***p<0.001 post-hoc Dunn’s. See Table S1. MN=motoneurons. Genotypes: (A-D) Control: *w; D42GAL4/+*. (A, C) *w; D42GAL4/UASkek6RFP. w; D42GAL4/UASAla; w; D42GAL4/UASkek6RFP UASAla*. (B, D) *D42GAL4/UASCaMKIIRNAi. w; D42GAL4/UASkek6RFP UASCaMKIIRNAi*.

Pre-synaptic CaMKII localizes to active zones [49,55], so next we asked whether CaMKII was required for the increased active zones caused by *kek-6* gain of function. CaMKII inhibition with Ala or knock-down with RNAi decreased active zones (Fig.S4A,B and Fig. 9B, Table S1) and over-expression of *activated CaMKII* in motoneurons increased active zones (Fig.S4A,C, Table S1). Remarkably, over-expression of *Ala* or *CaMKII-RNAi* together with *kek6*, rescued the increase in active zones caused by *kek6* gain of function (Fig. 9C,D, Table S1). These showed that CaMKII is required downstream of Kek-6 for active zone formation. Furthermore, in *kek6^−/−^* mutants and *kek6^−/−^ DNT2^−/−^* double mutants the levels of pCaMKII^T287^ decreased (Fig. 10A,Table S1), showing that Kek-6 is required for CaMKII activation at the NMJ. We found no significant effect of *DNT2* loss or gain of function on pCaMKII^T287^ levels at the NMJ. Importantly, over-expressing activated *CaMKII^T287D^* pre-synaptically rescued the phenotypes of decreased axonal length and reduced Ib boutons of *kek6^−/−^* (Fig. 10B) and *DNT2^−/−^* (Fig. 10C) single mutants, and *kek6^−/−^ DNT2^−/−^* double mutants (Fig. 10D) (Table S1). This means that the mutant phenotypes were caused, at least partly, by decreased CaMKII activation. Together, these data demonstrate that Kek-6 and DNT2 function in concert upstream of CaMKII to regulate NMJ size and active zones.

**Fig. 10.**
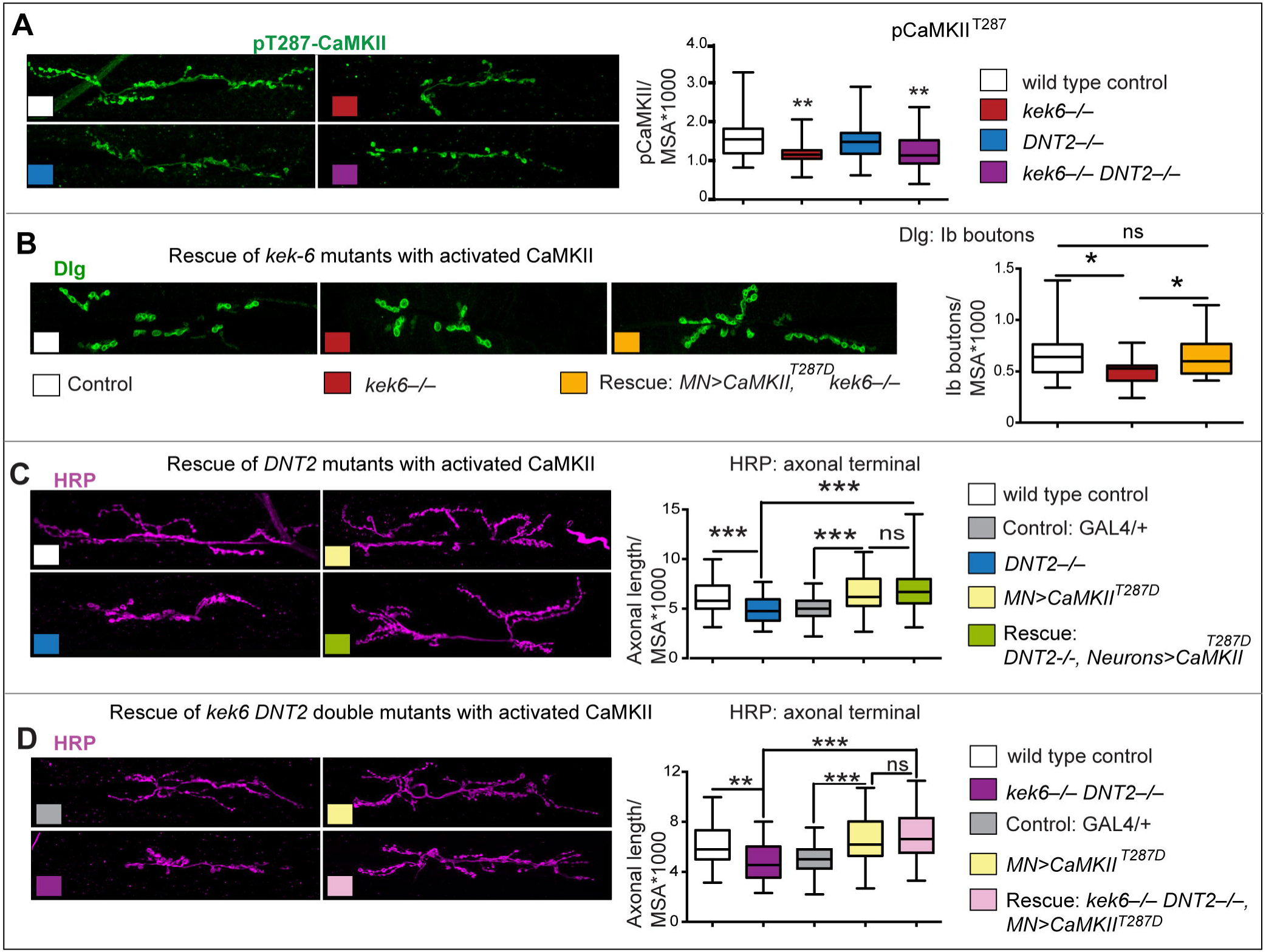
CaMKII functions downstream of Kek6 and DNT2 at the NMJ. Confocal images showing A3-4 muscle 6/7 NMJs, labeled as in Figure 9. (A) Reduced pCaMKII^T287^ levels in *kek-6* mutants, and *kek-6 DNT2* double mutants. Kurskal-Wallis p=0.0009, and post-hoc Dunn. (B-D) Over-expresison of activated *CaMKII^T287D^* in motoneurons rescued the NMJ phenotypes of (B) *kek-6* mutants, Kruskal-Wallis p=0.0081, post-hoc Dunn; (C) *DNT2* mutants, Kruskal-Wallis p=0.0001, post-hoc Dunn, and (D) *kek-6 DNT2* double mutants. Welch ANOVA p<0.000, post-hoc Games-Howell. *p<0.05, **p<0.01, ***p<0.001 see Table S1. Genotypes: MN=motoneurons, *D42GAL4;* Neurons: *elavGAL4*. Different neuronal drivers were used due to genetic constraints. Control: wild-type: *yw/*+; grey boxes: *D42GAL4/*+. (A) Mutant genotypes as in Fig. 4; Rescues: (B) *w;UASCaMKII^T287^/*+*; D42GAL4 kek6^34^/ Df*(*3R*)*6361*. (C) *w;UASCaMKII^T287D^/*+*; elavGAL4 Df*(*3L*)*6092/DNT2^37^*. (D) *w; UASCaMKII^T287D^/Toll-7GAL4; kek6^34^Df*(*3L*)*6092/Df*(*3R*)*6361 DNT2^37^*.

To test the functional link between Kek-6, DNT2 and VAP33A, we used genetic epistasis. Loss of *VAP33A* function in mutants caused a reduction in Ib boutons (Fig. 11A,B), and pre-synaptic over-expression of *VAP33A* increased Ib bouton number (Fig. 11A,D,E, Table S1), as previously reported[52]. These phenotypes were also shared with alterations in *kek6* and *DNT2* levels, consistent with common functions. Importantly, over-expression of *VAP33A* in motoneurons rescued bouton number in *kek-6^−/−^* (Fig. 11A,C) and *DNT2^−/−^*(Fig. 11A,D) single mutants (Table S1). Thus, VAP33A functions downstream of Kek-6 and DNT2. Interestingly, *kek-6^−/−^ DNT2^−/−^* double mutants rescued the increase in Ib boutons caused by *VAP33A* over-expression, restoring bouton number down to control levels (Fig. 11A,E, Table S1). This suggests that VAP33A may be required both for pre-synaptic vesicle release downstream of Kek6, and post-synaptic secretion of DNT2.

**Fig. 11.**
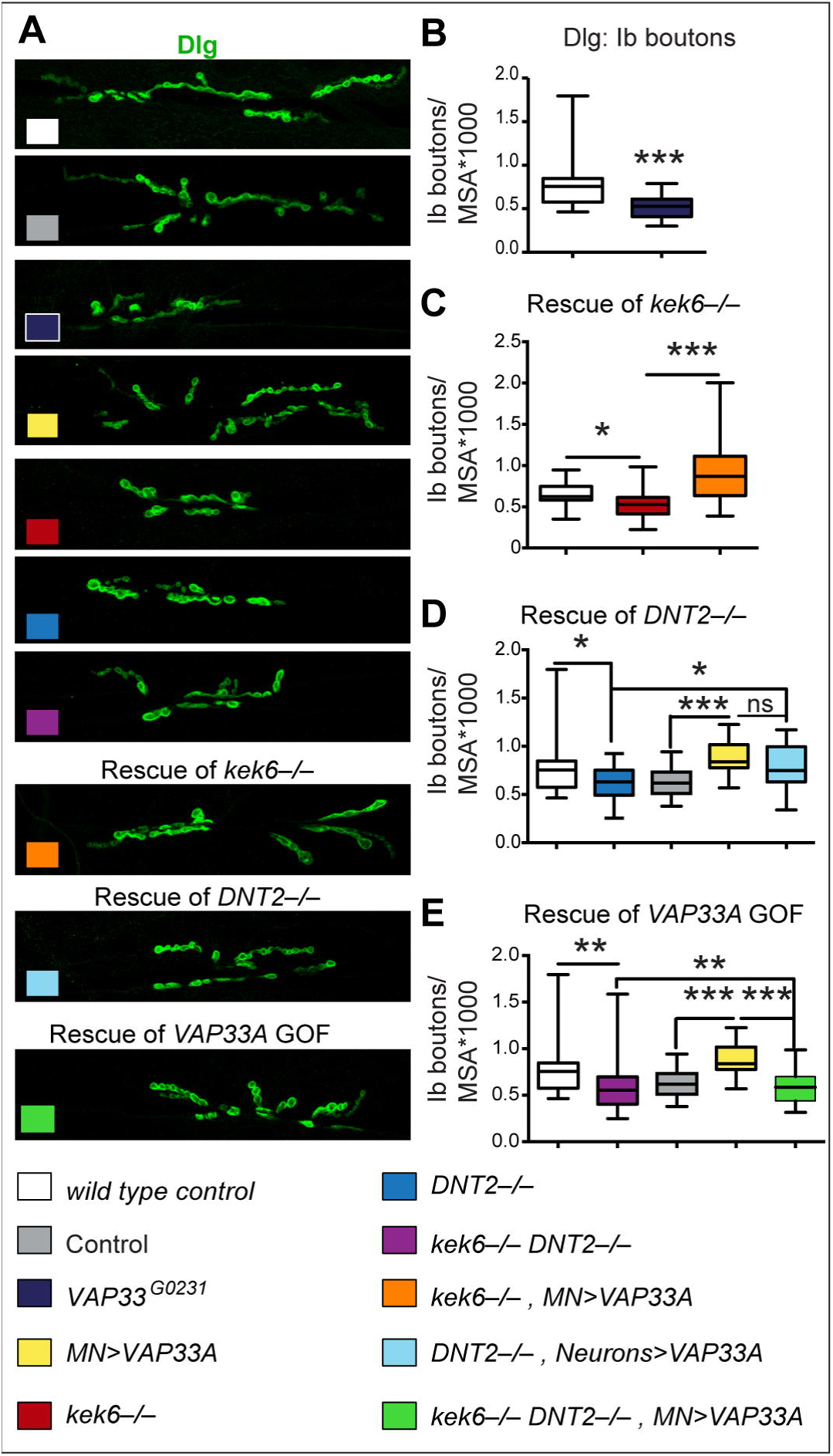
VAP33A functions downstream of Kek-6. (A) Confocal images of NMJs from segments A3-4, muscle 6/7. (B-E) Box-plot graphs. (A) *VAP33A^G0231^* mutants have reduced bouton number, Mann-Whitney U test ***p<0.001. (C,D) Pre-synaptic over-expression of *VAP33A* rescues bouton number in (C) *kek-6* mutants and (D) *DNT2* single mutants, Kruskal-Wallis p<0.0001 and *p<0.05, ***p<0.001 post-hoc Dunn for both. (E) *kek-6 DNT2* double mutants rescue the bouton number phenotype caused by *VAP33A* gain of function, Kruskal-Wallis p<0.0001 and **p<0.01, ***p<0.001 post-hoc Dunn. See Table S1. MN=motoneuron, *D42GAL4* (D) or *Toll-7GAL4* (E). Rescue genotypes: (C) *UASVAP33A; D42GAL4 kek6^34^/Df(3R)6361*. (D) UASVAP33A/+; *elavGAL4 Df(3L)6092/DNT2^37.^*. (E) *UASVAP33A/Toll-7GAL4; kek6^34^Df(3L)6092/Df(3R)6361 DNT2^37^*

To conclude, altogether these data show that CaMKII and VAP33A function downstream of DNT2 and Kek6 in motorneurons.

## DISCUSSION

Drosophila homologues of the Trk receptor family had long been sought. This was important to find fundamental principles linking structure and function in any brain. Here, we show that Kek-6 is a Trk-family receptor lacking a TyrK, for the neurotrophin ligand DNT2, regulating structural synaptic plasticity via CaMKII and VAP33A, in *Drosophila* (Fig. 12A,B).

**Fig.12.**
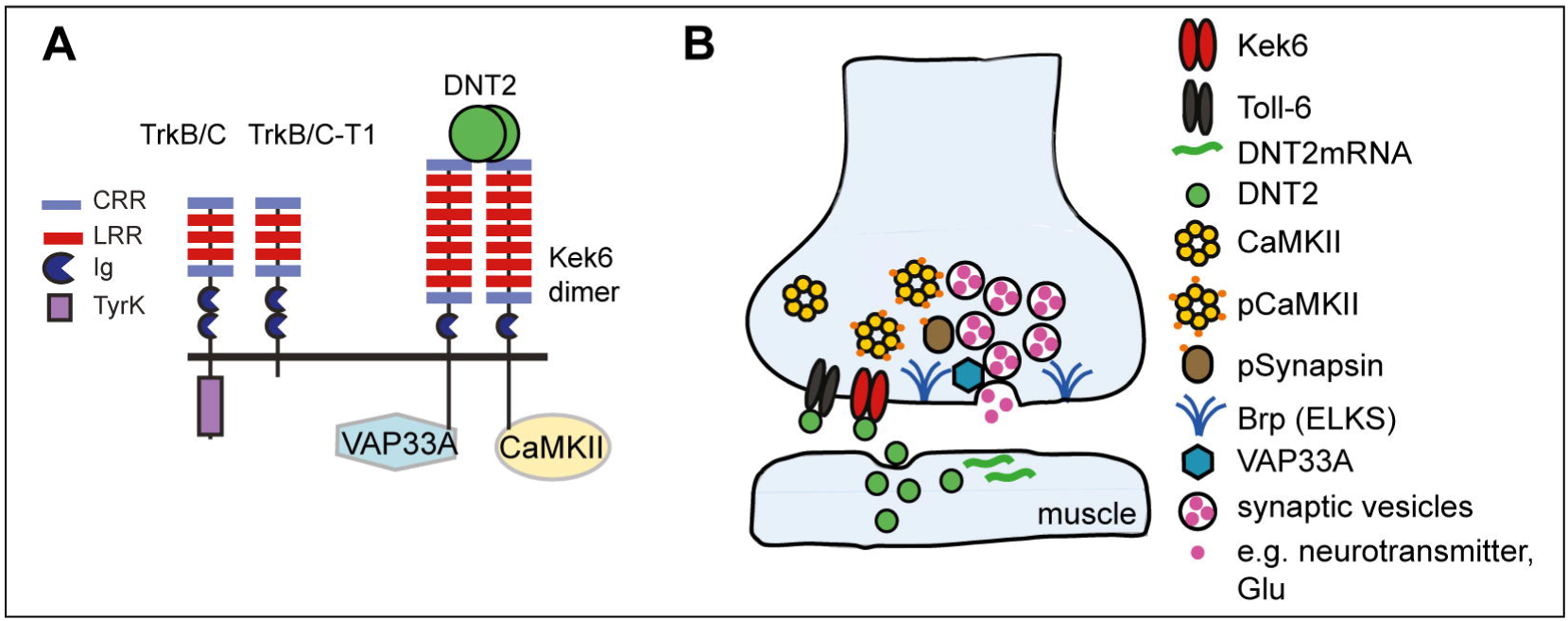
Retrograde DNT2 binds pre-synaptic Kek-6 activating CaMKII and regulating structural synaptic plasticity. (A) Illustration of Kek-6 compared to Trk isoforms, and showing that DNT2 binds Kek-6, which functions via CaMKII and VAP33A downstream. (B) Pre-synaptic motorneuron terminal at the NMJ: DNT2 is produced at the muscle and secreted, binds pre-synaptic Kek-6, functioning via CaMKII and VAP33A downstream to regulate NMJ growth and active zones. DNT2 also binds Toll-6 and interactions between Kek-6 and Toll-6 regulate NMJ structural homeostasis.

Starting with an unbiased approach, we showed that of all Drosophila LIGs Keks are the most likely DNT receptors in the CNS, and used two assays to demonstrate binding of Kek-6 to DNTs. Using in vivo analyses, we demonstrate that motoneurons express *kek-6*, and Kek-6 binds DNT2 which is produced by the muscle in post-synaptic boutons, and functions retrogradely. *kek-6* and *DNT2* share mutant phenotypes, *kek6 DNT2* double mutants have synergistic phenotypes, and *kek-6* over-expression can rescue mutant phenotypes of *kek6^−/−^* and *DNT2^−/−^* single single and *kek6^−/−^DNT2^−/−^* double mutants. Kek-6 is not a ligand sink or dominant co-receptor for Toll-6. Instead, Kek-6 and Toll-6 can interact physically and function cooperatively as DNT2 receptors. Kek-6 recruits CaMKII and VAP33A to a pre-snaptic downstream complex and induces CaMKII activation. Epistasis analysis demonstrated that Kek6 functions in motorneurons downstream of DNT2 and upstream of CaMKII and VAP33A to promote NMJ growth, and active zone formation. We conclude that DNT2 and its receptors Kek-6 and Toll-6 regulate synaptic structural plasticity and homeostasis.

Despite abundant evidence that retrograde signals and positive feedback loops regulate structural synaptic plasticity at the *Drosophila* glutamatergic NMJ, the identification of retrograde factors had been scarce [45,56], creating a void for the general understanding of structural synaptic plasticity. BDNF is a key retrograde ligand in mammals, and together with its receptor TrkB they form a positive feedback loop that reinforces synaptic function [1,4]. We have shown the neurotrophin DNT2 is a retrograde ligand for Kek6 (Fig. 12B). DNT2 is produced in muscle and over-expression of tagged DNT2 in muscle leads to its distribution in pre-synaptic boutons; over-expression of *DNT2* in muscle induces bouton, active zone formation and axonal terminal growth, pre-synaptically; *DNT2* mutants have smaller NMJs, and pre-synaptic over-expression of either *kek6, CaMKII* or *VAP33A* can rescue the *DNT2* loss of function phenotype; and conversely, *kek6* loss of function rescues the phenotype caused by *DNT2* over-expression in muscle. Thus, DNT2 is an evolutionarily conserved retrograde factor at the *Drosophila* synapse.

We demonstrate that *keks* are the *Drosophila Trk*-family homologues. Previous searches for Trk-family receptors in fruitflies had focused on the Tryrosine Kinase domain, and did not identify any bona fide candidates [14-16,18]. Subsequent proteomic analyses confirmed the absence of full-length Trk receptors with a conserved tyrosine kinase in fruitflies [19-21]. Since those earlier searches, Trks in mammals have been found to encode multiple isoforms, most of which lack the tyrosine kinase domain [6,57]. Human TrkB has 100 isoforms, which produce 36 different proteins, of which four are abundant, and the most abundant isoform in the adult human brain is truncated TrkB-T1, which lacks the tyrosine kinase [6,57]. Paradoxically, the function of TrkB-T1 in neurons in the mammalian brain is unknown. Our findings suggest that truncated Trk family receptors could have an evolutionarily conserved function regulating structural synaptic plasticity.

We also show that Kek-6 functions cooperatively rather than antagonistically with pre-synaptic Toll-6, a known receptor for DNT2, that is also required for synaptic structural plasticity [43]. LIG proteins and truncated Trk isoforms can function as ‘dominant negative’ co-receptors or ‘ligand sinks’. TrkB-T1 can form dimers with TrkB-FL and abrogate signaling, and bind BDNF rendering it unavailable to TrkB-FL [6,7]. This is thought to modulate kinase-signaling levels by TrkB [58]. Similarly, in *Drosophila*, Kek-5 is an inhibitor of BMP signaling [26] and Kek1 an inhibitor of EGFR signaling [59]. Thus, each Kek could function as a specific inhibitor of different signaling pathways. We have shown that Kek-6 and Toll-6 can form a receptor complex for DNT2, as DNT2, as well as binding Toll-6 [38], can also bind Kek-6, and Kek-6 and Toll-6 interact physically. However, our evidence shows that Kek6 does not function as an inhibitor of Toll-6, and instead Kek-6 and Toll-6 have distinct functions that positively cooperate to regulate NMJ structure and growth. All *kek-6^−/−^* and *Toll-6^−/−^* single mutants, and *kek-6^−/−^ Toll-6^−/−^* double mutants, have smaller NMJs, ruling out antagonistic functions. Toll-6 is required for active zone formation, and over-expression of *kek-6* also increases active zones, which means Kek-6 does not repress Toll-6. Interactions between DNT2, Kek-6 and Toll-6 are required for homeostatic compensation of active zones, as compensation fails in *kek6^−/−^ DNT2^−/−^* and *kek-6^−/−^ Toll-6^−/−^* double mutants. Our data suggest that Kek-6 and Toll-6 influence synaptic structure via alternative, independent pathways. Toll-6 functions in neurons via a canonical pathway to regulate NFκB and JNK [36]; Toll-8 and Toll-6 regulate NMJ plasticity also via an alternative dFOXO pathway[37,43]; and we show that Kek-6 functions via CaMKII and VAP33A to regulate synaptic structure independently of Toll-6 (Fig. 12B). With our assays, DNTs can bind both Tolls [38] and Keks (this work) promiscuously. The biological significance of this promiscuity is not understood, as ligand-receptor interactions will depend on the availability of each one in different contexts and times. Altogether, our findings strongly suggest that DNTs, Keks and Tolls constitute a novel molecular mechanism that regulates synaptic structural plasticity and homeostasis.

We do not yet know how CaMKII gets activated downstream of Kek-6. CaMKII activation and recruitment to active zones depends on the intracellular increase in Ca^2+^ levels, both through Ca^2+^ channels, and Ca^2+^ stores through the PLC and IP3 pathways [50,55]. In mammals, TrkB-T1 regulates Ca^2+^ levels in glia and in the heart, two contexts expressing high levels of TrkB-T1 and no TrkB-FL [60-62]. However, the mechanism by which TrkB-T1 raises Ca^2+^ is unknown [60,61], and whether this results in the activation of CaMKII is unclear or may be context dependent [60,62]. CaMKII activation in neurons could depend on the increase in Ca^2+^ with neuronal activity. A potential link of Kek-6 to neuronal activity was revealed by the increase in ghost boutons with *kek-6* gain of function. Furthermore, *kek-2* expression is modulated by neuronal activity, appropriate Kek2 levels at the NMJ are required for normal synaptic structure, and DNT2 can modulate the Na+/K+ ATPase [27,63]. Thus, Kek-6 could function via a PLC pathway or by influencing membrane channels, to increase Ca^2+^ levels and induce CaMKII activation.

The finding that Trk-like receptors can regulate neuronal plasticity independently of kinase signalling is very important. Activation of PLCγ by TrkB TyrK signaling is generally seen as a key mechanism of plasticity [1,5]. Yet CaMKII is necessary and sufficient for synaptic structural plasticity, LTP and long-term memory, and has wide functions in synaptic organization and homeostasis [49-51]. CaMKII can function as a frequency detector, and cause long lasting changes in synaptic strength, structure and brain plasticity [50]. Thus, the regulation of CaMKII and Ca^2+^ by truncated Trk-family receptors and Keks means that they could regulate structural synaptic plasticity independently of the canonical TyrK-dependent PLCγ pathway.

Keks are not identical to truncated Trk receptors and may carry out further functions that could be implemented via other routes in mammals. TrkB-T1 has a very short intracellular domain, whereas Keks have longer intracellular fragments including a PSD95/Dlg/ZO1 (PDZ) motif [22,24,25]. PDZ domains are involved in scaffolding, and assembly of post-synaptic complexes that regulate the size and strength of synapses [64]. Through its PDZ domain, Kek-6 might recruit synaptic partners and Ca^2+^ channels. Interestingly, truncated TrkC-T1 binds the PDZ-containing protein Tamalin to induce cellular protrusions via Rac[65]. Thus, it is imperative to explore further functional conservation or divergence of signaling mechanisms of Keks vs. truncated Trks.

To conclude, evolution may have resolved how to implement structural synaptic plasticity through distinct mechanisms in fruit-flies and humans, or perhaps common molecular principles are shared by both, even if not in every detail. We found that in fruitflies, truncated-Trk-like receptors encoded by the Keks bind neurotrophin ligands to regulate structural synaptic plasticity via CaMKII and VAP33A, and the receptor complex also includes Tolls. It is compelling to consider whether such a non-canonical mechanism of neuronal plasticity, downstream of neurotrophins and Trk receptors but independently of kinase signaling, may also operate in humans, as it could uncover novel mechanisms of brain function and brain disease.

## ACKNOWLEDGEMENTS

We thank: our lab for discussions; M.Landgraf, G. Tear, J.Hodge, S.Brogna, L.Griffith, N.Perrimon, B. Pfeiffer, M.Ramaswami for reagents and/or advice; the Bloomington and Kyoto Stock Centres for flies; Developmental Studies Hybridoma Bank for antibodies; *Drosophila* Genome Resource Center for clones. This work was funded by Wellcome Trust Project Grant 094175/Z/10/Z to A.H. and N.G., Wellcome Trust Equipment Grant 073228/Z/03/Z to A.H., MRC-Career Establishment Grant G0200140 to A.H., MRC PhD studentship to G.M., Marie Curie International Incoming Post-doctoral Fellowship to J.W., BBSRC PhD studentship to S.B. and Science Without Borders-CAPES PhD Studentship BEX 13380/13-3 to S.U.B. The authors declare that they have no conflict of interests.

## MATERIALS AND METHODS

### Genetics. Mutants

*Df(3R)ED6361* lacks the kek-6 locus (Kyoto Stock Centre), *Df(3L)6092, Df(3L)ED4342, DNT1^41^* and *DNT2^e03444^* are described in[35], *Df*(3L)BSC578 lacks the *Toll-6* locus (Bloomington). Mutant null alleles *kek6^34^* and *kek6^35^*, and *DNT2^37^*, were generated by FRT mediated recombination of the PiggyBac insertion lines [46]: for *Kek6: PBac{RB}kek6^e000907^* and *PBac{WH}kek6^f05733^;* for *DNT2: PBac{RB}spz5^e03444^* and *PBac{WH}Shab*^f05893.^. Mutants were selected using genetics and PCR, using primers as recommended in [46]. *kek6^34^* is a 41.9 kb deletion that just removes the coding region for *kek6*; DNT2^37^ is a 27 kb deletion that removes the ATG and first exon, most likely resulting in no protein production (Foildi et al, under review). **GAL4 lines:** *kek5GAL4* line *P{GawB}NP5933* (from BSC); *w;;elavGAL4* drives expression in all neurons (insertion on the third chromosome); *Toll-6-GAL4* was generated by RMCE from *MIMICToll-6^MIO2127^; w;;D42GAL4* and *w;Toll-7GAL4* drive expression in motoneurons and have been previously described[38]; *w; MhcRFP MhcGAL4* drives expression in muscle (BSC); *w; GMRGAL4* drives expression in retina. **UAS lines**: to drive expression of each of the *keks* UAS-lines were made as described below. *w;;UAS-DNT2-FL* expresses full length untagged DNT2; *w;;UASDNT2-CK* expresses the mature form of DNT2, i.e. signal peptide+cystine-knot domains; *w; UAS-DNT2-FL-GFP* expresses full-length DNT2 tagged at the COOH end, and were made as described below. *w; 10xUASmyr-Td-Tomato* is membrane tethered (gift of B. Pfeiffer); *w;UASCaMKII^287D^* expresses constitutively active CaMKII and *w;UASAla* expresses the CaMKII inhibitor (gifts of J. Hodge); *UASCaMKII-RNAi: y sc v; P{y[*+ *v*+, *TRiP.GL00237}attP2/TM3, Sb* (BSC); *y[1] w[*]; P{w[*+*mC]*=*UAS-FLAG.Vap-33-1.HA}2* (*BSC*). Double mutant lines were generated by conventional genetics. **MIMIC-GFP lines**: *y w; MIMICkek6^MI13953^* and has a GFP insertion in the coding region (BSC). All stocks were balanced over *SM6aTM6B* or *TM6B* to identify balancer chromosomes, and all were generated from a *yw* or *w* mutant background. In all figures, controls are: (1) F1 from *yw x Oregon;* (2) *w; GAL4/*+

### Molecular cloning

Full length cDNAs for *kek1, 2, 3, 4, 5* and *6* were obtained either from cDNA clones (*kek1, SD01674; kek2, NB7*), by PCR from cDNA libraries (gift of G. Tear; LD: *kek5;* GH: *kek3; kek6*), or by reverse transcription PCR from larvae (*kek4*), using primers designed for Gateway cloning into the pDONR plasmid: *kek1* forward: GGGGACAAGTTTGTAC AAA AAA GCA GGC TCA TCC AGG AAA **ATG** CAT ATC A and reverse: GGGGACCAC TTT GTA CAA GAA AGC TGG GTA GTC AGT TCT TGG TTT GGT TT; *kek2*: forward: GGGGACAAGTTTGTAC AAA AAA GCA GGC TCA **ATG** AGT GGT CTG CCA ATC T and reverse: GGGGACCAC TTT GTA CAA GAA AGC TGG GTA AAT GTC GCT GGT TTC CTG GC; *kek3*: forward: GGGGACAAGTTTGTAC AAA AAA GCA GGC TCA TAT GCG **ATG** GCA GCG GGA A and reverse: GGGGACCAC TTT GTA CAA GAA AGC TGG GTA GCT CTT GAA AAT ATC CTG TC; *kek4*: forward: GGGGACAAGTTTGTAC AAA AAA GCA GGC TCA CTA GAC CTT CCG TTC CTT, and reverse: GGGGACCAC TTT GTA CAA GAA AGC TGG GTA TAT TGA GAT ATC AAC ACC AG; *kek5*: forward: GGGGACAAGTTTGTAC AAA AAA GCA GGC TAG CTA GAC GCA GAC TTA GAG and reverse: GGGGACCAC TTT GTA CAA GAA AGC TGG GTA GAC CTC GGT GCC ATC CTC GC; *kek6*: forward: GGGGACAAGTTTGTAC AAA AAA GCA GGC TCA **ATG** CAT CGC AGC ATG GAT C and reverse: GGGGACCAC TTT GTA CAA GAA AGC TGG GTA GAG CGA CAC GAA CTC GCC AG. *CaMKII* and *VAP33A* full-length cDNAs were cloned using the Gateway System first into *pDONR*, and then into *pAct5-attR-HA*.

For expression in flies under UAS/GAL4 control, cDNAs were subcloned to a Gateway *pUASt-attR-mRFP* destination vector for conventional transgenesis, injected by BestGene (www.thebestgene.com). For expression in S2 cells, they were subcloned into tagged *pAct* destination vectors *pAct5c-attR-mCFP, pAct5c-attR-FLAG* and *pAct5c-attR-HA*. Constructs used for expression of S2 cells of DNT1 and 2, Toll-6 and 7 were as previously described[38]. Chimaeric Kek-Toll-6 receptors were generated after analyzing the domain composition of the proteins using ProSite (ExPASy), PFAM, SMART, TMHMM and TMPred algorithms and PubMed data. Primers were designed to amplify the sequences that encode the extracellular and transmembrane domains of Kek3–6, including 15 amino acids C-terminal to the Kek transmembrane region, and the intracellular domain of Toll6, at halfway between the transmembrane region and the TIR domain. A unique enzyme site (BamHI or EcoRI) was included in the designed primers to join the *kek* and *Toll-6* sequences, and attB sites were introduced in the primers at the 5′ and 3′ ends of the chimaeric insert for Gateway cloning into *pAct5c-attR-3xHA* destination vector. Unfortunately, we were not able to generate *kek1-Toll-6* and *kek2-Toll-6* chimaeric receptor constructs. To clone chimaeric *kek3*,*4*,*5*,*6-Toll-6* receptor constructs, reverse primers at the juxtamembrane of *keks* were as follows: *kek3-Toll-6:_CGAT*-GAATTC-AGGTACAGAGTTCCAGAGAC; *kek4-Toll-6*: CGCG-GAATTC-TTGCAAATAAGTGTGCTGGC; *kek5-Toll-6*: CTAT-GGATCC-GCTCATCATGGTGGTGTCCT; *kek6-Toll-6*: GTAT-GAATTC-ACGCCGGCCTTGTTGGCATG. *Toll-6* primers from chimaeras were: Forward primers from juxtamembraneCATG-GAATTC-AACTTCTGCTACAAGTCACC (compatible with *kek3, kek4* and *kek6* chimaeras) or CATG-GGATCC-AACTTCTGCTACAAGTCACC (compatible with *kek5* chimaera) and reverse from C terminus: GGGGACCAC TTT GTA CAA GAA AGC TGG GTC CGC CCA CAG GTT CTT CTG CT.

Un-tagged full-length DNT2 was cloned into the pUASt-attB vector by conventional cloning, DNT2-full length (DNT2-FL) was PCR-amplified from cDNA libraries, and cloned into pUASt-attB for ΦC31 transgenesis [66]. UAS-DNT2-FL-GFP was cloned by Gateway cloning into pUAS-GW-GFP, followed by conventional transgenesis.

### Luciferase signaling in S2 cells

For Dif signaling assays using chimaeric kek-Toll-6 receptors, S2 cells stably transfected with *drosomycin-luciferase* were maintained at 27^0^C in air in Insect-Xpress medium (Lonza) supplemented with penicillin/streptomycin/L-glutamine mix (Lonza) and 10% foetal bovine serum (Lonza). 1ml suspended cells were passaged every two days into 4ml fresh medium. 3×10^6^ cells in 2ml media were seeded per well of a 6-well plate 24 hours prior to transfection. Per well of experiment, 250μl serum-free media, 3μl TransIT-2020 (Mirus), 2μg of HA tagged chimaeric receptors *kek-toll6-HA* plus 1μg of *pAct-renilla-luciferase* were incubated at room temperature for 30 minutes, supplemented with 350μl serum-free media and added to aspirated cells. After 4 hours, transfection mixture was removed and 2ml supplemented medium added. All experiments were conducted 48 hours after transfection; for imaging membrane targeting of Kek-Toll6-HA protein S2 cells were starved for 6h prior to fixation.

For signaling assays, S2 cells were stimulated with 50nM/well of purified baculovirus DNT2 protein (generated as previously described[38]) and Dif signalling quantified by luminescence. DNT2 was added 48 hours after transfection and luminescence quantified 24 hours after DNT stimulation. Transfected and stimulated cells were pelleted from single wells, resuspended in 400μl media and separated into three 50μl aliquots in an opaque 96-well plate. 40μl of Firefly Luciferase Substrate (Dual-Glo Luciferase Assay System; Promega) was added per 50μl aliquot, incubated for 10 minutes, and luminescence measured using a Mithras LB 940 Multimode Microplate Reader (Berthold). 40μl Stop & Glo substrate (Dual-Glo Luciferase Assay System; Promega) was added to quench Dif signal and activate Renilla Luciferase. Renilla luminescence data was used to normalize Firefly Luciferase data.

### Co-immunoprecipitations

Coimmunoprecipitations were carried out as previously described[38], after transfecting standard S2 cells with the following constructs: (1) Co-IP Kek6-DNT2: *pAct5C-Pro-TEV6HisV5-DNT2-CK* and *pAct5C-Kek6-3xHA;* (2) Co-IP Kek6-DNT2 or Kek6-DNT1: *pAct5C-Kek6-3xHA, pAct5C-Pro-TEV6HisV5-DNT1-CK-CTD, pAct5C-Pro-TEV6HisV5-DNT2-CK;* (3) Co-IP Kek5-DNT2: *pAct5C-Kek5-3xFlag* and *pAct5C-Pro-TEV6HisV5-DNT2-CK*. (4) Co-IP kek6-Toll-6: *pAct5C-Kek6-HA* and *pAct5C-Toll-6^CY^-3xFlag, pAct5C-Toll-7^CY^-3xFlag*¨ (5) Co-IP Kek6-CaMKII: *pAct5-CaMKII-HA* and *pAct-kek-6-FLAG*. (6) Co-IP kek6-VAP33A: *pAct5-Vap33A-HA* and *pAct-kek-6-FLAG*. 48h after transfection cells were harvested and washed in PBS. Final cell pellets were lysed in 600 μL NP-40 buffer (50 mM Tris-HCl pH:8.0, 150 mM NaCl, 1% Igepal-630) supplemented with protease inhibitor cocktail (Pierce). For V5 and HA immuno-precipitations,500 μL of lysates from single or co-transfected cells were incubated with 1 μg of mouse anti-V5 or 1 μg of mouse anti-HA antibodies overnight at 4 °C, then the lysate plus antibody mixtures were supplemented with 25 μL of protein-A/G megnetic beads (Pierce) and incubated for 1 h at room temperature. For Flag immunopreciptations, lysates were incubated with anti-Flag antibody conjugated magnetic beads (Sigma-Aldrich) overnight at 4 °C. In both cases, beads were washed thoroughly in NP-40 buffer and/or PBS. Proteins were eluted in 40 μL of 2x Laemmli-buffer and analysed by Western blot following standard procedures.

### Pull-down assays and proteomics

S2 cells were transfected with 4 μg *pAct5C-Kek6-3xFlag* expression construct. For controls mock-transfected cells (i.e. no construct) were used. After transfection cells were processed as described above for co-immunoprecipitation. 600 μL of cell lysates were incubated with 20 μL of anti-Flag conjugated magnetic beads overnight at 4 °C. Beads were washed thoroughly in PBS, then proteins were eluted in 40 μL 2x Laemmli-buffer for 10 min at room temperature. Eluted proteins were loaded onto 10% polyacryalamide gels. Alternatively, proteins were not eluted after overnight incubation with anti-Flag beads, but they were re-incubated with whole fly (OregonR) head lysates. Here, 60 heads were lysed in 600 μL of NP-40 buffer. After overnight incubation at 4 °C, proteins were eluted and analysed as for S2 cell lysate proteins. Thus, eluted proteins were loaded onto 10% polyacryalamide gels, and the gels were stained with Coomassie Blue and cut into several pieces. Gel pieces were subjected to in-gel digestion with trypsin using a standard protocol. Tryptic peptides were analysed by LC-MS/MS using Ultimate 3000 HPLC coupled to a LTQ Orbitrap Velos ETD mass-spectrometer. Peptide separation, mass spectrometric analysis and database search were carried out as specified at the University of Birmingham Proteomics Facility. Candidate binding proteins were identified on these criteria: (1) Proteins were accepted only if they were identified with at least two high confident peptides. (2) Mock-transfected controls and Kek6-3xFlag samples were compared, and only proteins identified from Kek6-3xFlag lysates or heads, but absent form controls, were considered as possible interacting partners for Kek6.

### Western blotting

Western blot was carried out following standard procedures using antibodies listed here: mouse anti-V5 (1:5000, Invitrogen), mouse anti-HA (1:2000, Roche), chicken anti-HA (1:2000, Aves), rabbit anti-Flag (1:2000, Sigma-Aldrich), HRP-conjugated anti-mouse IgG (1:5000-1:10000, Vector), HRP-conjugated anti-chicken IgG (1:5000, Jackson Immunoresearch), HRP-conjugated anti-rabbit IgG (1:1000-1:5000, Vector), mouse dCaMKII (Cosmo CAC-TNL-001-CAM 1:1000), rabbit α-p-CaMKIIα (Thr 286) (Santa Cruz sc-12886-R 1:1000), mouse Tubulin DM1A (Abcam ab7291 1:10000).

### NMJ dissections and preparation

NMJ preparations were carried out according to [67]. For GFP stainings in *MhcGAL4*>*UAS-DNT2-FL-GFP*, L3 larvae were placed in agar plates (2%) and left at 29°C for 90 minutes to potentiate the NMJ before dissections [68]. A hundred and twenty female flies and 50 males were placed in a cage with a removable agar and grape juice plate. On the second day, plates were changed for new ones in the morning and evening, and discarded. On the third day, the plates were changed every 1h 30min and discarded. On the fourth day, the plates were changed every 1h 30min and kept in a 25°C incubator for larvae collection. The next day, hatched L1 larvae from each plate were transferred to a vial, and exactly 40 larvae were placed in each vial. When larvae reached L3 stage (5 days after egg laying), they were dissected in low calcium saline as previously described[67], and fixed for 10 minutes in Bouin’s solution (HT10132 SIGMA). Samples were washed 6 times for 10 minutes in PBT (0.1% of Triton in 1M PBS) to remove fixative solution and kept overnight in blocking solution (10% normal goat serum in 0.1% Triton in PBS 1M). Primary antibodies were incubated overnight at 4°C and samples were washed the following day 8 times for 10 minutes in PBT. Secondary antibodies were incubated for 2h at room temperature and samples were washed 8 times for 10 minutes in PBT. Samples were mounted in Vectashield anti-bleacing medium (Vector Labs) in No.0 coverslips. NMJs were analysed for muscle 6/7 only; segments A3 and A4 were analysed in each larva.

### Immuno-stainings and in situ hybridisations

Antibody stainings in embryos and S2 cells were carried out following standard procedures, using the following primary antibodies at the indicated dilutions: mouse anti-FasII at 1:5 (ID4, Developmental Studies Hybridoma Bank, Iowa); rabbit anti-GFP 1:1000 (Molecular Probes); mouse anti-Dlg 1:20 (4F3, Developmental Studies Hybridoma Bank, Iowa); rabbit anti-HRP 1:250 (Jackson Immunoresearch); mouse anti-Brp 1:100 (nc82, Developmental Studies Hybridoma Bank, Iowa); rabbit anti-p-CaMKII^T287^ 1:150 (Santa Cruz), detects phosphorylation at Thr286 (T287 in *Drosophila*), constitutively active form; mouse plain anti-Synapsin at 1:25 (DSHB 3C11). Secondary antibodies were: anti-guinea pig-Alexa 488 at 1:250 (Molecular Probes); biotynilated anti-mouse at 1:300 (Jackson Labs) followed by the ABC Elite Kit (Vector Labs); biotynilated antiguinea pig at 1:300 (Jackson Labs) followed by Streptavidin-Alexa-488 at 1:400 (Molecular Probes); anti-rabbit-Alexa 488 at 1:250 (Molecular Probes); anti-mouse-Alexa 488 1:250 (Molecular Probes); anti-rabbit-Alexa 647 at 1:250 (Molecular Probes); anti-mouse-Alexa 647 1:250 (Molecular Probes).

In situ hybridizations were carried out following standard procedures, using antisense mRNA probes from 5’ linearised plasmids and transcribed as follows: *lambik*: a 551 nucleotides fragment was cloned into pDONR with primers: forward GGGGACAAGTTTGTAC AAA AAA GCA GGC TAG AAA CTA CGC ATG AGC CTG and reverse GGGGACCAC TTT GTA CAA GAA AGC TGG GTA CCG CTC AAA TGT CCA CTG T; then linearised with HpaI and transcribed with T7 RNA polymerase. *CG15744*: a 569 nucleotides fragment was cloned into pDONR with primers: forward GGGGACAAGTTTGTAC AAA AAA GCA GGC TGG ATT GGA TAG CCT TGG TGA and reverse GGGGACCAC TTT GTA CAA GAA AGC TGG GTT TCG CTT CCA TCT CCA TCT C (linearised with HpaI, transcribed with T7); *CG16974*: a 548 nucleotides fragment was cloned into pDONR with primers: forward GGGGACAAGTTTGTAC AAA AAA GCA GGC TTA TAT GAA TCC CGA AGG CGC and reverse GGGGACCAC TTT GTA CAA GAA AGC TGG GTT TGG GGG GAG TAG ATG GTA A_(linearised with HPA1, transcribed with T7); *kek1* (*SD01674*+*pOT2* cDNA clone; linearised with EcoRI, transcribed with SP6 RNA polymerase); *kek2* (*NB7*+*pNB40;* HindIII; T7); *kek3* (HpaI; T7); *kek4* (*GH27420*+*pOT2;* EcoRI; SP6); *kek6* (in *pDONR*, HpaI; T7); *CG15744* (in pDNOR, HpaI; T7); *CG16974* (in pDNOR, HpaI; T7); *lambik* (in pDONR, HpaI; T7). Colorimetric reaction was using Alkaline Phosphatase conjugated anti-DIG conjugated.

### Microscopy and Imaging

Wide-field Nomarski optics images were taken with a Zeiss Axioplan microscope, 63x lens and JVC 3CCD camera and Image Grabber graffics card (Neotech) and a Zeiss AxioCam HRc camera and Zen software. Laser scanning confocal microscopy was carried out using a Leica SP2 AOBS inverted confocal microscope, 40x lens, at 1024 x 1024 resolution, and a Zeiss LSM710 inverted laser scanning confocal microscope with a 40x lens, 1024 x 1024 resolution, 0.25μm step for anti-Brp, and 0.5 μm step for the rest. Images were compiled using ImageJ, Adobe Photoshop and Illustrator.

For NMJ data, all images provided in the figures are projections from Z stacks; all the quantitative anlyses were carried out in the raw stacks, not in projections. Muscle surface area was measured from bright field images using ImageJ. Anti-HRP was used to measure total terminal axonal length using ImageJ. Boutons were visualized with anti-Dlg, and total boutons as well as separately Is and Ib boutons, were counted manually with the aid of the ImageJ Cell Counter plug-in. Automatic quantification of active zones labeled with anti-Brp, and anti-pCaMKII^T287^, were carried out in 3D throughout the stacks of images using the DeadEasy Synapse ImageJ plug-in that we previously validated [39], and anti-Synapsin was analysed with a slightly modified version. DeadEasy Synapse first reduces the Poisson noise, characteristic of confocal microscopy images, of each slice in the stack using a median filter. Subsequently, in order to separate signal (e.g. Brp+ active zones) from the background, images are segmented. Since the intensity of the staining varied within each image and from image to image, the maximum entropy threshold method [69] was used to find a local optimal threshold value for each pixel. For this, we used a square window of size 15x15, sufficient to find the local optimal threshold around each pixel in an image. In this way, each pixel is considered part of an active zone if the value of the pixel is higher than the local threshold, otherwise assigned to the background. Since this method is computationally expensive and very low intensity pixels correspond to background, it was possible to reduce the computation time by assigning pixels whose intensity was lower or equal than 20 directly as part of the background and only applying the thresholding method to pixels whose intensity was higher than 20. Finally the volume of the active zones is measured. This method worked just as accurately with Brp, pCaMKIIT^287^ and Synapsin stainings. Data were normalized to muscle surface area or axonal length.

### Locomotion assays

L3 wandering larvae were placed one at a time on an agar plate (2%) and left it crawl for 40 seconds. Larvae were filmed crawling across the agar plate and then discarded. Plates were cleared before placing another larva. At least 50 larvae were filmed per genotype, and the test was always done in the morning. Films of 400 frames per larva were analysed using FlyTracker software developed in our lab to obtain the trajectory and speed, as previously described[38].

### Statistical analysis

Data were analysed in SPSS Statistics 21 (IBM) and GraphPad Prism 6. Confidence interval was 95% (p<0.05). Categorical data were tested using X^2^, and a Bonferroni correction was applied for multiple comparisons. Continuous data were first tested for normality by determining the kurtosis and skewness, and a Levene’s Test was applied to test for homogeneity of variance. Data were considered not normally distributed if absolute kurtosis and skewness values for each genotype were greater than 1.96 x standard error of kurtosis/skewness. Variance of the populations of different samples were considered unequal if Levene’s test for homogeneity of variance gave a p value of <0.05. If samples were normally distributed and variances were equal, Student-t tests were applied for 2- sample type graphs, and One-Way ANOVA was used to compare means from >2 samples. When data were normally distributed, but variances were unequal, Welch ANOVA was used instead. Multiple comparison corrections to normal data were applied using a post-hoc Dunnett test of comparisons to a control, Games-Howell or Bonferroni post-hoc comparing all samples. Non-parametric continuous data were compared using a Mann-Whitney U-test when 2 sample types were being analysed, and a Kruskal-Wallis test for >2 samples, and multiple comparison corrections were applied using a post-hoc Dunn test. See Table S1 for all statistical sample sizes, applied tests and p values.

## SUPPORTING INFORMATION

**Fig. S1 Kek-6 is expressed is not expressed in glia.** Confocal images of the VNC neuropile of third instar Kek-6^MIMIC-GFP^ larvae, stained with anti-GFP and the pan-glial nuclear marker anti-Repo. Repo does not colocalise with GFP in any cells, arrows point to examples of Repo+, GFP-negative cells.

**Fig. S2 Altered *kek-6* function affects motoraxon targeting at embryonic NMJ.** The motoneuron marker FasII reveals motoraxon targeting phenotypes at muscle 6,7,12,13 in stage 17 embryos, in *kek-6* mutants and upon over-expression of *kek-6* in all neurons (with *elavGAL4*). Arrows indicate stereotypic projections in wild-type, and mistargeting in other genotypes. Chi-square p<0.0001 and ***p<0.001 Boferroni corrections, see Table S1.

**Fig. S3 *kek-6* over-expression induces ghost boutons.** (A,B) Over-expression of *kek-6* in motorneurons (MN) with *D42GAL4* did not affect bouton number (Dlg, Mann-Whitney U-test not significant). (C-E) Over-expression of *kek-6* induced pre-synaptic ghost boutons lacking a post-synaptic component (arrows: HRP+, presynaptic and Dlg-negative, post-synaptic), (D) higher magnification; (E) quantification. Both bouton number and area increased, albeit not significantly. Mann-Whitney U-tests. See Table S1. *Genotypes*: Control: *D42GAL4/*+. : *kek-6^−/−^: kek6^34^/Df*(*3R*)*6361; MN*>*kek6: D42GAL4*>*UAS-kek6-RFP* (with one or two copies of *UAS-kek-6*).

**Fig. S4 Loss and gain of CaMKII function affect the NMJ.** (A) Confocal images of muscle 6/7 NMJs, in A3-4, labeled with anti-HRP for the pre-synaptic terminal, anti-Brp for active zones, anti-Dlg for post-synaptic boutons and anti-pCaMKII^T287D^ for the constitutively active form. (B,C) Quantification. (A,B) Inhibiting CaMKII function with Ala in motoneurons (*D42GAL4*>*UASAla*) decreased NMJ terminal axonal length (HRP,t-test), and active zones. Student t-tests, p<0.005, **p<0.01. (A,C) Pre-synaptic over-expression of constitutively active CaMKII (*D42GAL4*>*UASCaMKII^T287D^*) increased pCaMKII levels (pCaMKII^T287D^, Student t-test **p<0.001), axonal length (HRP, Student t-test **p<0.001), Ib bouton number (Dlg, Mann-Whitney U-test ***p<0.001), and active zones (Brp, Mann-Whitney U-test **p<0.01). See Table S1.

**Table S1 Statistical Analysis Details.** For all genotypes, sample sizes, tests and p values, see this table.

**Table S2 Kek6 pull-down from S2 cells.** List of S2 cell lysate proteins binding Kek6-Flag, but not mock-transfection controls, and detected with more than one peptide.

**Table S3 Kek6 pull down from adult fly heads.** List of adult fly head proteins binding Kek6-Flag, but not mock controls, and detected with more than one peptide.

